# Immunomodulatory Contribution of Mast Cells to the Regenerative Biomaterial Microenvironment

**DOI:** 10.1101/2020.07.31.231852

**Authors:** Raymond M. Wang, Joshua M. Mesfin, Jessica L. Ungerleider, Yu Kawakami, Yuko Kawakami, Toshiaki Kawakami, Karen L. Christman

## Abstract

Bioactive immunomodulatory biomaterials have shown promise for influencing the immune response to promote tissue repair and regeneration. Macrophages and T cells have been associated with this response; however, other immune cell types have been traditionally overlooked. In this study, we investigated the role of mast cells in the regulation of the immune response to decellularized biomaterial scaffolds using a subcutaneous implant model. In mast cell-deficient mice, there was dysregulation of the expected M1 to M2 macrophage transition typically induced by the biomaterial scaffold. Polarization progression deviated in a sex specific manner with an early transition to an M2 profile in female mice, while the male response was unable to properly transition past a pro-inflammatory M1 state. Both were reversed with adoptive mast cell transfer. Further investigation of the later stage immune response in male mice determined a greater sustained pro-inflammatory gene expression profile including the IL-1 cytokine family, IL-6, alarmins, and chemokines. These results highlight mast cells as another important cell type that influences the immune response to pro-regenerative biomaterials.

## Introduction

The immune response is known to have a necessary role in tissue homeostasis, remodeling, and repair, and our understanding of the involved immune populations and their phenotypic state improves our capacity to treat adverse disease conditions. Mast cells are tissue resident granulocytes classically associated in host defense, pro-inflammatory responses and various immune disorders^1^. The granules of mast cells are known to contain a variety of immune mediators demonstrated to contribute to pathological responses such as allergic reactions^2^, inflammatory cell recruitment^3^, and fibrosis^4^. While mast cells are also known to secrete a plethora of pro-remodeling/anti-inflammatory cytokines that are relevant for tissue repair, their role in tissue regenerative therapies has not been studied.

Biomaterial scaffolds that influence the immune response are emerging as a viable clinical strategy for tissue repair and regeneration^5^. This immunomodulatory influence associated with improved tissue outcomes has often been characterized as an early pro-inflammatory response that transitions to a pro-remodeling response at later timepoints^6–8^. Although this induced response has been rigorously evaluated with macrophage^7,9^ and T cell populations^10,11^, potential contribution of mast cells to the immunomodulatory effect of naturally derived biomaterials has been largely ignored and under investigated^12^. *In vitro* culture of human mast cell lines has demonstrated that interaction in the decellularized extracellular matrix (ECM) microenvironment promotes mast cell differentiation, maturation, and viability compared to collagen controls^13^. However, *in vivo* studies have mainly just evaluated shifts in mast cell numbers associated with biomaterial deployment, though their general low numbers in tissue make it difficult to determine conclusive results from this approach^14,15^. Early studies with synthetic materials assessing their functional role have mainly produced results characteristic of a classical mast cell phenotype highlighting contributions to early acute inflammatory cell infiltration^3^, and fibrous capsule formation^16^. Thus, there has been limited investigation on the influence mast cells have on the immune response to naturally derived biomaterials *in vivo*.

From a general tissue disease perspective, several studies have similarly emphasized the detrimental role of mast cells to tissue remodeling outcomes^17–19^. However, research has also produced confounding results^20–23^ or evidence of their beneficial contributions to select cases of native tissue repair or inflammatory resolution^21,24,25^, suggesting a greater functional complexity and potential to be utilized in therapeutic design. Given the recent clinical and rapidly expanding pre-clinical use of decellularized ECM biomaterials, which are known to induce pro-remodeling and pro-regenerative effects based on immunomodulatory mechanisms in macrophages and T cells^9–11^, we chose to evaluate this class of biomaterial scaffold in mast cell-deficient mice compared to wild-type and mast cell engrafted mice. Mast cell deficient Kit^W-sh^ mice were utilized as they are highly deficient in mast cells across various tissues and in the skin by 10-12 weeks while maintaining relatively normal levels of a broad range of lymphocyte and myeloid lineages^26^. For our investigation of the interaction of mast cells with a biomaterial scaffold, we used an injectable hydrogel form of myocardial matrix, which was previously shown to be immunomodulatory^8,27^ and recently progressed through a Phase I clinical trial in post-myocardial infarction patients^28^. Based on recent evidence suggesting that mast cells can have beneficial influences on native tissue remodeling outcomes, we hypothesized that mast cells play a role in the regulation of decellularized ECM biomaterial scaffold-induced immunomodulatory responses, and demonstrated in this investigation their critical regulatory contribution to the pro-inflammatory to pro-remodeling immunomodulatory transition, which deviates in a sex specific manner.

## Results

### Mast Cell Recruitment and Degranulation *In Vivo* in Response to ECM biomaterial

Limited studies have investigated the interaction of infiltrating mast cells and naturally derived biomaterials in the *in vivo* setting aside from the observation of increased recruitment^15,27^. In this study, decellularized ECM hydrogels were subcutaneously injected into the dorsal region of C57BL6/J wild-type mice and Kit^W-sh^ mast cell-deficient mice, and isolated at various timepoints. A subcutaneous implant model was chosen since we and others have previously shown it can recapitulate the pro-remodeling immune cell polarization response of decellularized ECM materials seen in disease or injury models^8,29^. As mast cells have demonstrated sex divergent responses *in vitro^30^*, *in vivo^31^*, and mast cell associated diseases clinically^32,33^, both male and female mice were initially assessed to determine whether biomaterial responses were consistent across sexes. Isolated subcutaneous injections showed mast cell presence at low cellular density as early as day 1 post-injection. Mast cells were non-degranulated or minimally degranulated based on toluidine blue staining (Fig. 1a). As expected, deficient mice showed no mast cell presence in the material or throughout the skin, confirming a lack of localized mast cell presence or recruitment (Fig. 1b). Greater inward infiltration of non-degranulated mast cells was observed for ECM scaffold isolated at 3 days post-injection (Supplementary Fig. 1a, b). Minimal to no fibrous capsule formation was observed around subcutaneous injections as also seen in a previous study^8^. Comparison with subcutaneous injection of commercial milled urinary bladder matrix showed similar recruitment of non-degranulated mast cells bordering the material at day 3 post-injection, demonstrating that these observations were not specific to the hydrogel form of decellularized ECM (Supplementary Fig. 1c, d). Similar to a previous study utilizing human mast cell lines to study the influence of decellularized ECM on mast cell viability^13^, interaction of differentiated bone marrow derived FcεRI^+^c-kit^+^ mast cells (Supplementary Fig. 2) with the ECM hydrogel material in solution at non-gelling concentrations in comparison to collagen and media only controls showed higher alamarBlue™ readings, thus, confirming matrix interaction enhanced differentiated mast cell viability (Supplementary Fig. 3). These results supported that mast cell interaction with ECM biomaterials promoted an alternative phenotypic state instead of the classically activated degranulated phenotype seen in allergic and foreign body type responses.

**Fig. 1:**
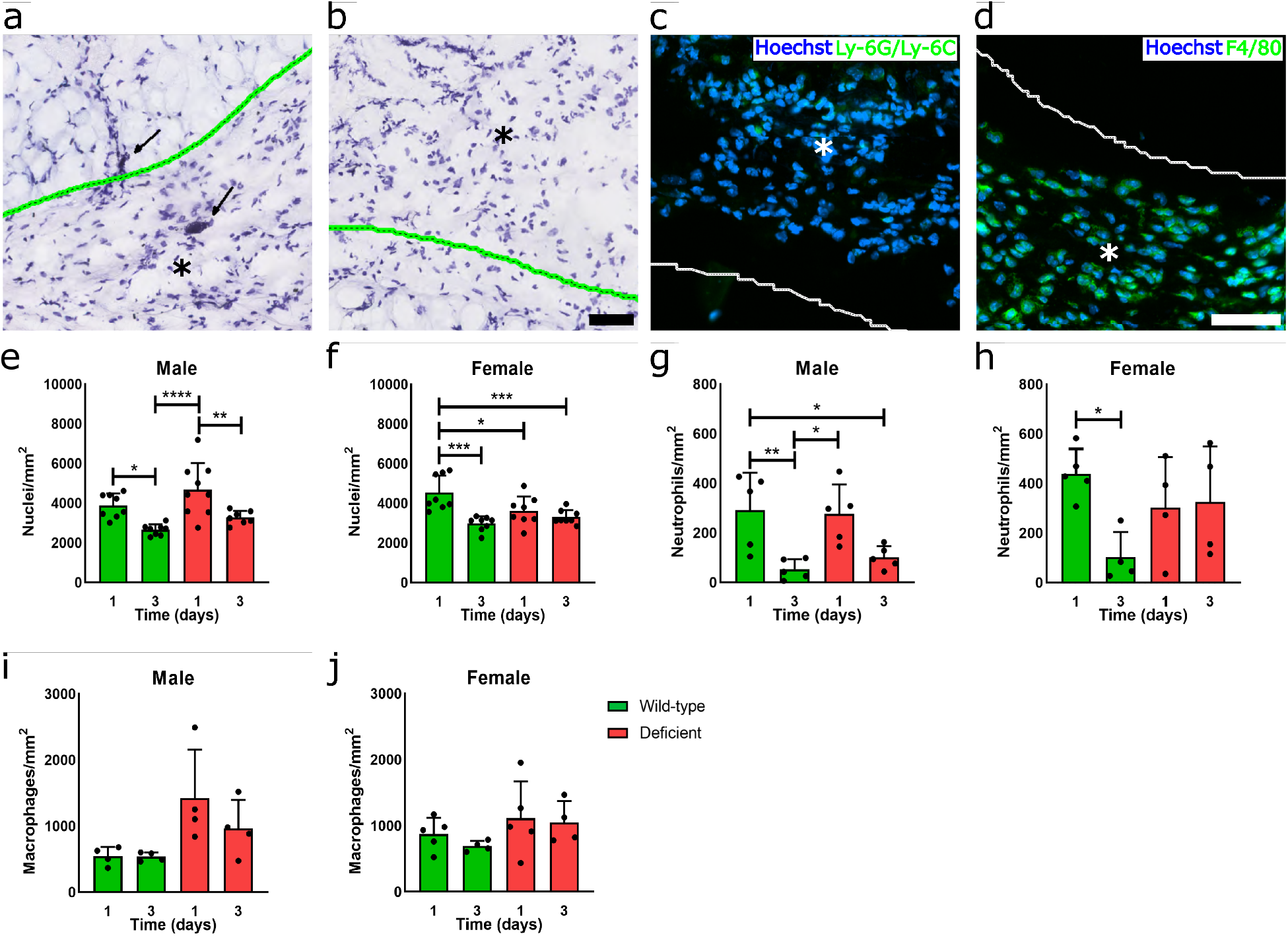
ECM scaffold promotes early immune cell infiltration with or without mast cells. Representative images of toluidine blue staining of ECM scaffold (green outlined with asterisk where material is present) and neighboring dermal tissue in wild-type **(a)** and mast cell-deficient **(b)** mice at day 1 post-injection. Mast cells can be found as early as day 1 showing lack of or minimal degranulation (black arrow). Representative fluorescent images of injected biomaterial (white outlined with asterisk) with Ly-6G/Ly-6C^+^ neutrophil **(c)** and F4/80^+^ pan-macrophage staining **(d)** in green, and Hoechst nuclei counterstain in blue. Quantification of total nuclei **(e, f)**, neutrophil **(g, h)** and macrophage **(i, j)** density at one and three days post-injection for male and female wild-type (green) and mast cell deficient (red) mice. n = 7-9 per group for total nuclei; n = 4-5 per group for immune cell quantification. Scale bar of 50 μm in toluidine blue **(a, b)** and fluorescent images **(c, d)**. (\**p* < 0.05, \*\**p* < 0.01, \*\*\**p* < 0.001 and \*\*\*\**p* < 0.0001).

### Similar or Increased Leukocyte Recruitment Despite Lack of Mast Cell Contribution

One known physiological function of mast cells is the release of histamine and chemokines promoting vascular permeability and recruitment of immune cells, such as neutrophils^34,35^ and macrophages^36^, to sites of inflammation or in response to foreign agents. In mast cell deficient mice, this recruitment was mitigated in the response to synthetic biomaterial implants^3^. However, properties of pro-regenerative ECM biomaterials have been demonstrated to drive and rely on the infiltration of a variety of immune cell populations into the scaffold for promoting immunomodulatory responses^37^. Thus, we investigated whether an ECM biomaterial could elicit immune cell infiltration in mast cell deficient mice compared to recruitment in wild-type C57BL6/J mice. Total cellular infiltration into the ECM hydrogel scaffold between deficient and wild-type mice was quantified along with staining for Ly-6G/Ly-6C^+^ neutrophil (Fig. 1c) and F4/80^+^ macrophage populations (Fig. 1d). In male mice, similar nuclei density was observed between mouse models that was initially higher at day 1 and significantly decreased at day 3 with no significant difference between models at matching timepoints (Fig. 1e). In contrast, a higher total cell infiltration was observed in female mice at day 1 for wild-type compared to deficient mice, though this difference was lost by day 3 (Fig. 1f). Assessment of neutrophils notably showed a similar high to low density change pattern between day 1 to 3, likely from the characteristic early neutrophil-driven immune response that, again, did not differentiate between mouse models (Fig. 1g). Female mice also showed no significant difference between models at either timepoint, though there was a higher average neutrophil density maintained at the day 3 timepoint (Fig. 1h). Additionally, staining for total macrophages showed no significance between either sex or model, though averages were generally higher in the deficient model (Fig. 1e-j). Overall, these results demonstrated that early immune cell infiltration from ECM biomaterial implantation does not depend on mast cell contribution.

### Induced Macrophage Polarization is Dysregulated and Rescued based on Mast Cell Presence

Another mast cell function is the release of a varied secretome with components that are known to regulate various phases of the immune response. As ECM scaffold injection allowed for early cellular infiltration in mast cell-deficient mice similar to wild-type mice despite the lack of mast cell contribution (Fig. 1), it was investigated whether the lack of this regulatory contributor influenced macrophage polarization and the expected pro-inflammatory to pro-remodeling immunomodulatory transition elicited from an ECM biomaterial^6,7,38^. Timepoints up to 11 days post-injection were evaluated since earlier studies have shown that macrophage polarization during this window is representative of long-term outcomes^39–41^. Additionally, the ECM hydrogel undergoes complete degradation within 2-3 weeks following injection, which complicates evaluation at later timepoints.

A comprehensive flow cytometry panel (Supplementary Fig. 4-5), including comparison of M2/M1 macrophage polarization by count ratio determined a lack of expected polarization shift towards being M2 dominant^8,11^ in male mast cell-deficient mice at day 11 post-injection compared to a significant shift in wild-type mice (Fig. 2a). To confirm that this was mast cell specific, mast cell-engrafted mice were also similarly assessed where adoptive transfer of bone marrow derived mast cells differentiated *in vitro* (Supplementary Fig. 2) were implanted subcutaneously into deficient mice at least 5 weeks preceding biomaterial injection to reconstitute the local resident mast cell population as previously described^26,42^. Mast cell engraftment was confirmed at this later timepoint by toluidine blue staining showing presence of mast cells in dermal tissue and extracted ECM scaffold injections (Supplementary Fig. 6). These engrafted mice showed a restoration of the M2 shift similar to wild-type mice, which was also significant compared to the deficient group (Fig. 2a). Assessing the individual percentages macrophages showed similar percentages of M2 and M21 macrophages for the wild-type and deficient mice while a higher relative number of M1 macrophages were present in deficient male mice at the day 11 (Supplementary Fig. 7). Notably, addition of mast cells in the engrafted mice had a significantly greater percentage of M2 macrophages compared to deficient mice (Supplementary Fig. 7c). In contrast, female mice showed an M2 dominant presence at the early day 3 timepoint (Fig. 2b), which arose from a higher relative percentage of M2 macrophages and not from differences in M1 macrophage percentages (Supplementary Fig. 7d, f). Evaluation of other immune cell types using flow cytometry including T cells (Supplementary Fig. 8), B-cells (Supplementary Fig. 9), dendritic cells (Supplementary Fig. 10), and mast cells (Supplementary Fig. 11) showed limited differences between the deficient, wild-type, and engrafted mouse models with the exception of the expected absence of mast cells in the deficient mice. Therefore, these results support that mast cells have limited impact on the infiltration of immune subpopulations in response to ECM biomaterials, but do contribute to the relative polarization of macrophage phenotypes that deviate in a sex specific manner.

**Fig. 2:**
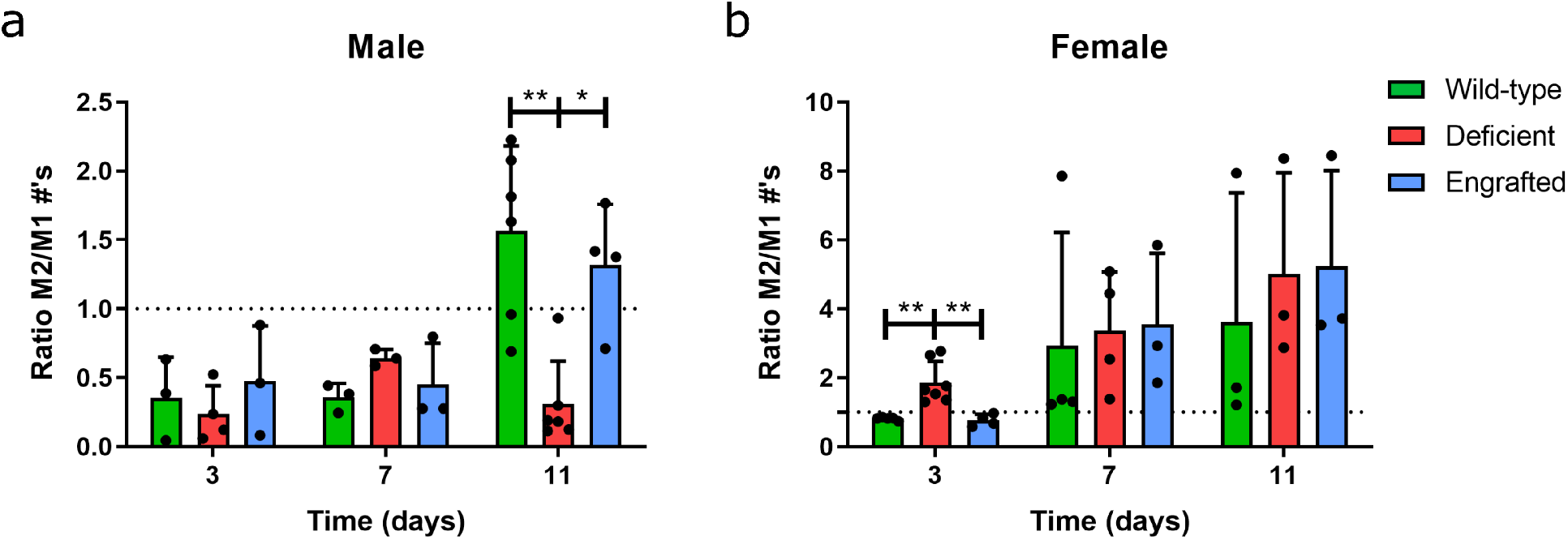
Transition from a pro-inflammatory to pro-remodeling macrophage is dysregulated in mast cell deficient mice in a sex specific manner. Flow cytometry quantification for macrophages polarization based on a ratio of CD206^+^ M2 versus CD86^+^ M1 macrophage counts was assessed at three, seven, and eleven days post-injection between wild-type (green), mast cell-deficient (red), and mast cell-engrafted (blue) male **(a)** and female mice **(b)**. n = 3-6 per group. (\**p* < 0.05, \*\**p* < 0.01).

Finally, to contextualize the degree of altered macrophage polarization observed in the mast cell deficient mice, we compared the response of our decellularized ECM biomaterial in the mast cell deficient model to an injectable, pro-inflammatory control consisting of a similarly processed non-decellularized material (NDM) that still contained cellular debris. In a separate cohort of mice, H&E staining of injected NDM confirmed that an immune response characteristic of immune rejection still occurs in mast cell deficient mice (Fig.3a), which was distinct from the decellularized ECM material (Fig. 3b). Since the NDM did not gel and was not distinguishably maintained up to the day 11 timepoint, we used a different validated macrophage polarization assessment based on Arg1/Nos2^8,43^ gene expression ratio where the M1-to-M2 transition was expected by day 7 post-injection^8^. At day 7 post-injection, a trending or significantly greater M2 dominant response for decellularized materials based on Arg1/Nos2 was determined for the wild-type and engrafted response in both sexes. In contrast, the deficient response was non-significant with a similar average between the decellularized and non-decellularized materials in both sexes (Fig. 3c,d). These results further demonstrate that mast cell deficiency leads to alteration in the M1-to-M2 transition that occurs with ECM biomaterials.

**Fig. 3:**
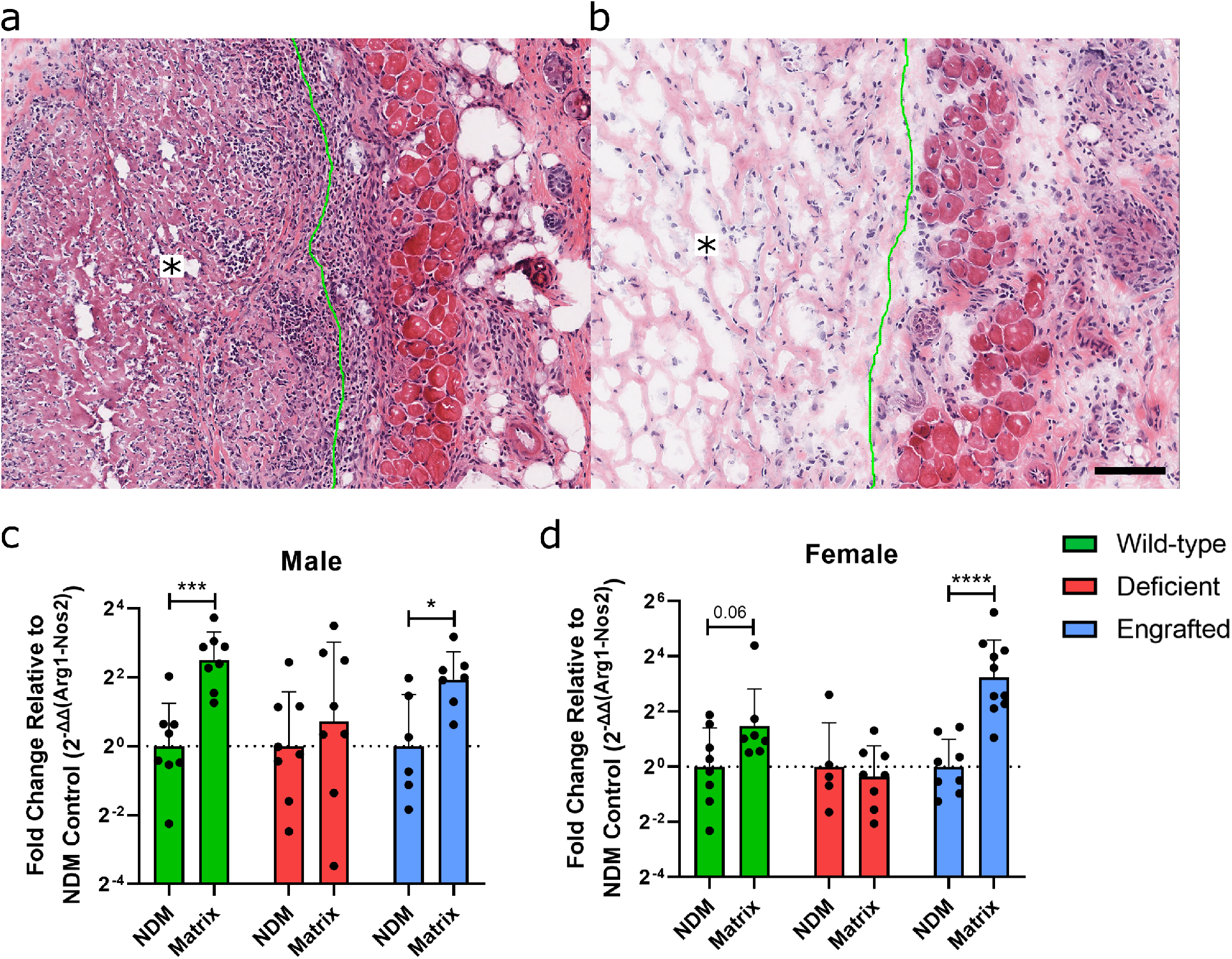
Comparison of mast cell influence in response to non-decellularized and decellularized ECM scaffolds shows mast cell deficiency leads to alteration in the M1-to-M2 transition with decellularized ECM. Representative hematoxylin and eosin staining in 11-13 week old deficient mice following dorsal subcutaneous injection of **(a)** non-decellularized and **(b)** decellularized ECM scaffold at 7 days post-injection (material outlined in green with asterisk on side where material is present). Scale bar is 100 μm. Gene expression ratio at day 7 post-injection of M2/M1 macrophage markers Arg1 relative to Nos2 in **(c)** male and **(d)** female mice. Decellularized ECM scaffold in wild-type (green), mast cell-deficient (red), and mast cell-engrafted (blue) groups were compared by relative gene expression with 2-^ΔΔct^ normalized to non-decellularized (NDM) control per group. n = 6-10 per group. (Value displayed for trend *p* ≤ 0.1, \**p* < 0.05, \*\*\**p* < 0.001 and \*\*\*\**p* < 0.0001).

### Pro-inflammatory Cytokine Expression is Similarly Dysregulated and Rescued based on Mast Cell Presence

Gene expression of several pro-inflammatory and pro-remodeling/anti-inflammatory markers (Table S1) were also assessed to confirm whether additional immune regulatory components were dysregulated based on mast cell absence. At day 3 post-injection, only a significant increase in anti-inflammatory Il38 in male deficient versus wild-type mice was found (Supplementary Fig. 12a). At day 11 when significant shifts based on flow cytometry analysis were observed, significant differences in the IL-1 cytokine family, including increases in pro-inflammatory Il1b and Il33, and decreases in Il38, was determined (Supplementary Fig. 12b). Further comparison with mast cell engrafted male mice showed these altered expression profiles were reversed to levels similar to wild-type expression for Il1b and Il33 (Fig. 4a, b), and for the gene expression ratio of IL-1 receptor antagonist, Il1rn, relative to Il1b (Fig. 4c). Expression of Cd206 was also significantly upregulated in engrafted mice compared to wild-type controls (Fig. 4d). Assessment for female mice at day 3 when dysregulation of macrophage polarization was observed by flow cytometry (Fig. 2b) determined only a significant upregulation of M1 marker Nos2 in deficient mice compared to wild-type that was reversed in engrafted mice (Fig. 4e and Supplementary Fig. 13). While this might seem to conflict with flow results showing a M2 dominant phenotype at this timepoint, Nos2 can be released by other cell populations and has been shown to be influenced by sex specific female hormones^44^. Assaying for gene expression of estrogen receptors, Esr1 and Esr2 (Table S2), determined a greater average expression for both receptors in the deficient mouse response compared to other groups with a significantly greater Esr2 expression for the deficient versus wild-type response (Fig. 4f, g). Thus, Nos2 expression could potentially be a less representative M1 marker in female mice due to hormonal influence that is exasperated with the additional loss of mast cells as an immune regulatory contributor. Overall, these results demonstrate the influence of mast cells on the expression of the IL-1 cytokine family in the ECM biomaterial response in male mice. In contrast, the response in female mice seems mainly a result of shifts in macrophage response; however, potential influence from female hormones makes these results difficult to interpret.

**Fig. 4:**
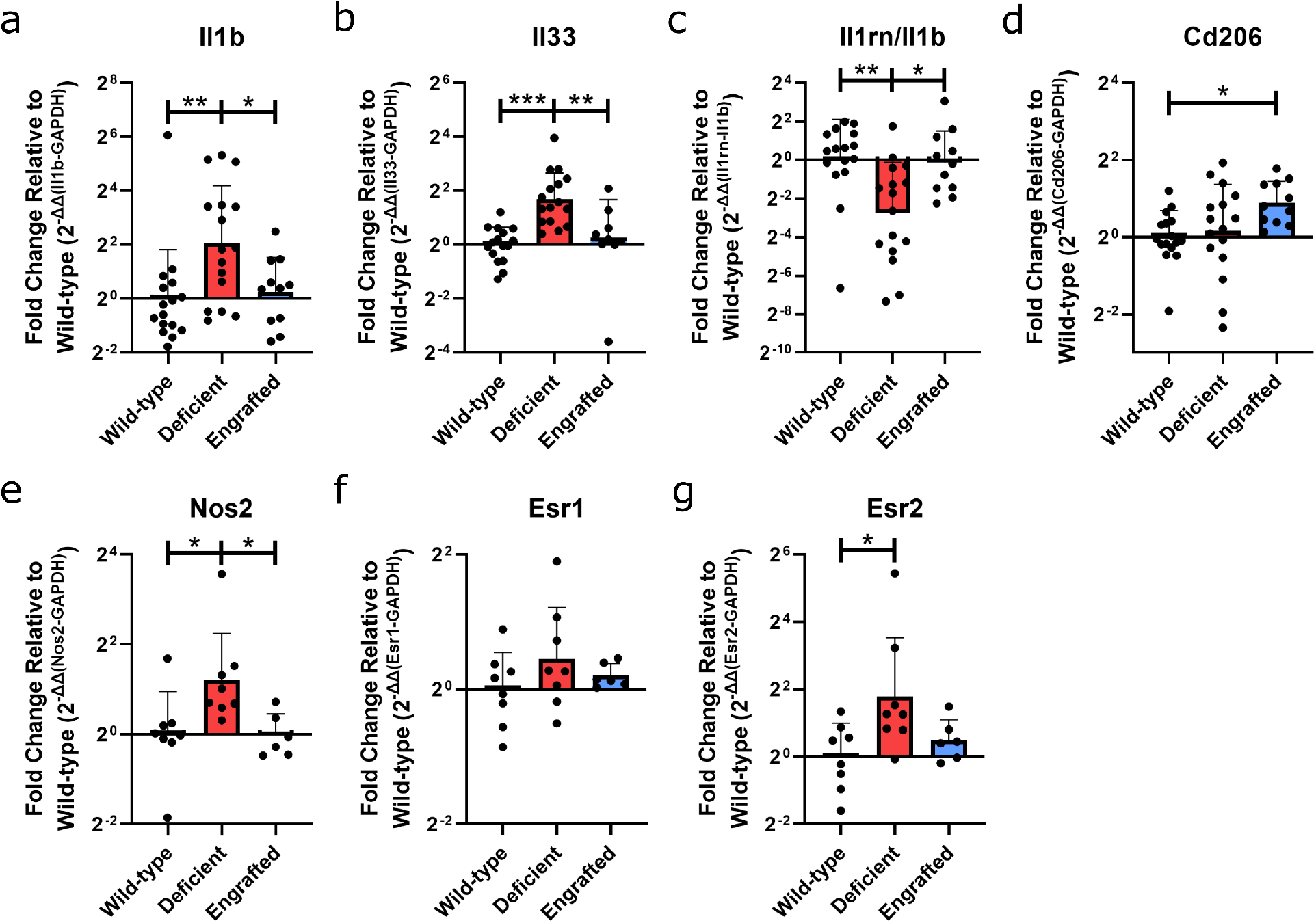
Immune marker profiling and hormone receptor gene expression demonstrating differentiation expression based on mast cell presence. qRT-PCR gene expression in deficient and engrafted normalized to wild-type expression by 2^-ΔΔct^ at day 11 in male mice for **(a)** Il1b, and **(b)** Il33 relative to housekeeping gene, Gapdh. **(c)** Gene expression ratio of receptor antagonist, Il1rn, relative to Il1b. **(d)** Gene expression of Cd206 relative to housekeeping gene, Gapdh. qRT-PCR gene expression at day 3 in female mice measured for **(e)** Nos2, **(f)** Esr1, and **(g)** Esr2 relative to Gapdh. n = 11-16 per group for male samples, n = 6-8 per group for female samples. (\**p* < 0.05, \*\**p* < 0.01, and \*\*\**p* < 0.001).

### Differential Pro-Inflammatory Expression Profile based on Mast Cell Presence

As mast cells are known to secrete and influence a broad spectrum of immune responses, a more comprehensive multiplex analysis was carried out through a Nanostring panel (raw data and analysis results provided). Only day 11 male mice samples were assessed by Nanostring (Supplementary Fig. 14) since the lack of pro-inflammatory resolution was observed in male deficient mice at this timepoint and because of the potential confounding variables in the female mice immune response from significant hormone contributions (Fig. 4). PCA analysis of the top 50% of genes based on variance showed that the majority of wild-type and engrafted samples clustered in separate groups along PC1 with some samples dispersed between the two clusters. Mast cell deficient samples were more distinct along PC2 and not as tightly clustered, potentially conveying increased variance due to lack of regulatory influence from mast cells (Supplementary Fig. 15). Generalized multi-group differential expression analysis of the deficient compared to the wild-type and engrafted responses determined differentially upregulated genes for a host of pro-inflammatory associated cytokines (Il1b, Il33, Il6, Tnf), chemokine and associated receptors (Ccl2, Ccl3, Ccl4, Ccl7, Cxcl1, Cxcl3, Cxcr2, Tslp), alarmins (S100a8, S100a9), enzymes (Ptgs2), and regulatory receptors (Clec4e, Il1rl1, Trem1). Additionally, upregulation of immune associated adhesion proteins (Muc1, Sele, Sell), defensins (Defb14), complement pathway proteases (Masp1), and growth factors (Ppbp) were determined (Fig. 5a). Notably, mast cell marker, Fcer1a, was also significantly decreased for the mast cell deficient group compared to wild-type and engrafted in both multi-group and pairwise comparisons, but not between the engrafted and wild-type samples (Fig. 5b, c and Supplementary Fig. 16), thus, further confirming successful mast cell engraftment.

**Fig. 5:**
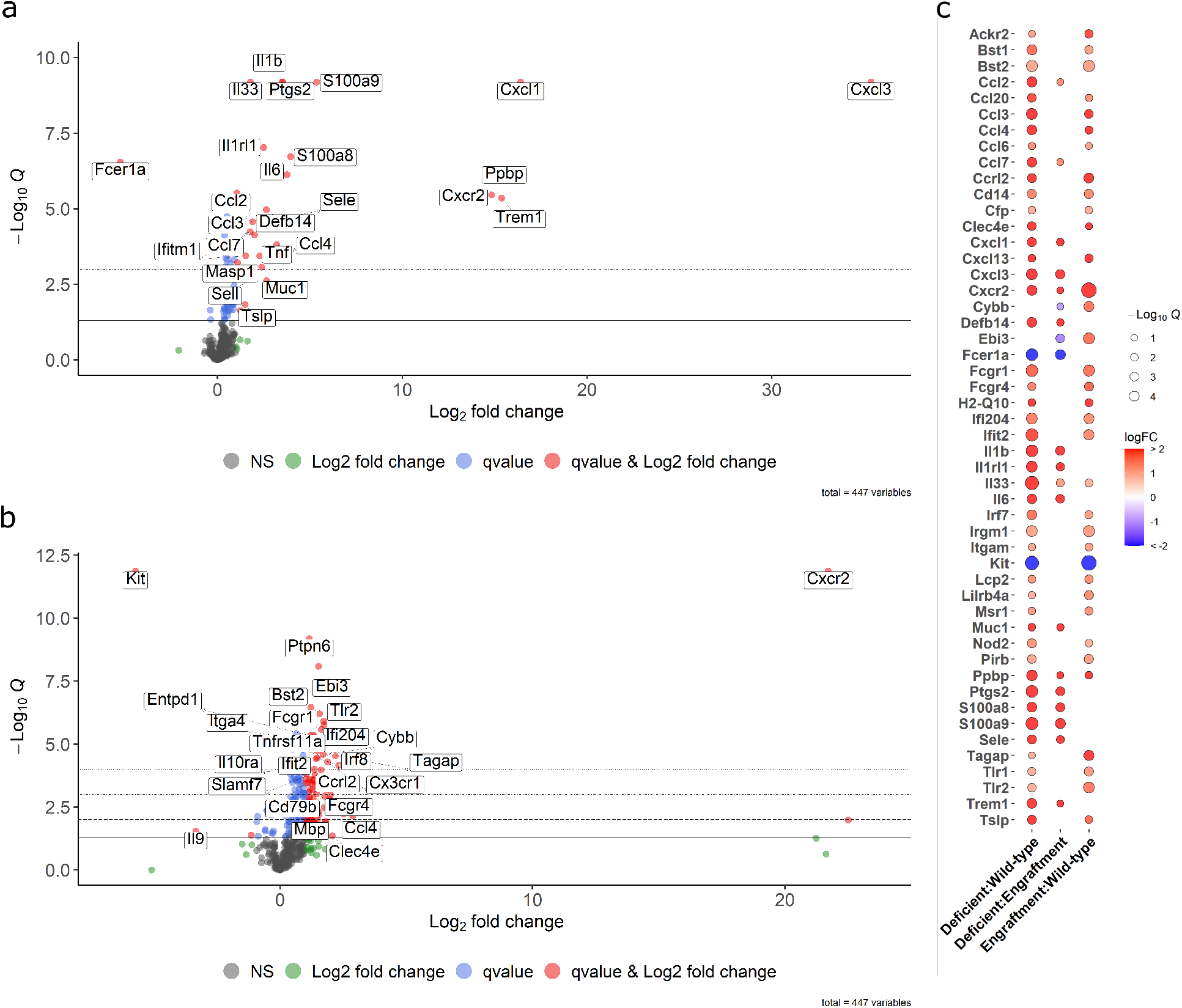
Mast cell-deficient male mice maintain late pro-inflammatory gene expression profile in response to ECM biomaterial implantation. Volcano plot of the top 25 differentially expressed genes with **(a)** mast cell deficient in a multi-group comparison against both wild-type and mast cell engrafted groups, and **(b)** pairwise comparison between engrafted to wild-type samples. Dot colors indicate *q* < 0.05 (blue, red), absolute fold change greater than 2 (green, red) or non-significant (black). Dashed lines indicating different significance thresholds (\**q* < 0.05, \*\**q* < 0.01, \*\*\**q* < 0.001 and \*\*\*\**q* < 0.0001). **(c)** Bubble plot showing significant differentially expressed genes consistent across at least two pairwise analyses amongst the wild-type, deficient, and engrafted groups. Red to blue color scale displays log2 fold change that was up- or downregulated, respectively, and circle size displays significance as −log10 of q-values. Results from n = 8 per group.

Pairwise analysis showed a similar pro-inflammatory response profile comparing deficient with wild-type alone (Ccl2, Ccl3, Ccl7, Cxcl1, Cxcl3, Il1b, Il1rl1, Il33, Il6, Ptgs2, S100a8, S100a9, Sele, Tnf, Trem1) (Supplementary Fig. 16a). Although the total number of differentially expressed genes between mast cell deficient and engrafted samples was more limited, notably these predominantly consisted of the various previously highlighted pro-inflammatory markers (Ccl2, Cxcl1, Cxcl3, Il1b, Il1rl1, Il33, Il6, Ptgs2, S100a8, S100a9, Sele, Trem1), conveying an altered inflammatory profile specifically related to the loss or presence of the localized mast cell response (Supplementary Fig. 16b). Furthermore, pairwise differential expression of the engrafted to wild-type mice highlighted upregulation of cellular signaling components (Cd79b, Entpd1, Mbp, Ptpn6), membrane oxidases (Cybb), transcription factors (Pou2f2), immune cell and immunoglobulin related regulatory receptors (Cd79b, Fcgr1, Fcgr4, Il10ra, Itga4, Slamf7, Tlr2, Tnfrsf11a), regulatory transcription factors (Irf8), immune cell activating proteins (Tagap), IFN-induced proteins (Ifi204, Ifit2), cytokine components (Ebi3), chemokines and associated receptors (Ccl4, Cx3cr1, Cxcr2), and downregulation of Il9 cytokine, which regulates cell growth and apoptosis (Fig. 5b). All together, these results convey an alternatively regulated immune profile in the engrafted versus wild-type response compared to the distinctly greater pro-inflammatory profile of the mast cell deficient response.

Finally, specifically investigating significant differentially regulated genes consistent across at least two pairwise comparisons amongst the three sample groups confirmed matching upregulation of previously highlighed pro-inflammatory markers (Ccl2, Ccl7, Cxcl1, Cxcl3, Cxcr2, Il1b, Il1rl1, Il33, Il6, Ptgs2, S100a8, S100a9, Trem1), defensins (Defb14), and adhesion proteins (Muc1, Sele) in deficient versus either wild-type or engrafted responses. In contrast, the majority of these pro-inflammatory markers were not similarly upregulated between engrafted and wild-type samples. Instead, genes related to cell growth and signaling factors (Bst1, Bst2, Ppbp), immune regulatory receptors (Ackr2, Irf7, Lilrb4, Pirb, Tlr1, Tlr2), chemokines and associated regulatory receptors (Ccl2, Ccl3, Ccl4, Ccl6, Ccl20, Ccrl2, Cxcl13, Cxcr2, Tslp), MHC class I antigen presentation and signaling (H2-Q10, Pirb), immune cell activating proteins (Lcp2, Nod2, Tagap), integrins (Itgam), cytokine components (Ebi3), macrophage scavenger receptors (Msr1), IFN-induced proteins (Ifi204, Ifit2), complement pathway regulators (Cfp), regulatory GTPases (Irgm1), and myeloid marker (Cd14) were upregulated (Fig. 5c). Considering the distinctly different gene profile between the deficient and engrafted response, particularly for pro-inflammatory cytokines, these results support that local mast cell reconstitution alone can alter the immunomodulatory biomaterial response.

### Enrichment of Pro-Inflammatory Pathways in the Absence of Mast Cells

Gene set enrichment analysis was performed to evaluate trends that were consistent (Fig. 6a) or divergent (Fig. 6b) amongst different group comparisons for the deficient versus both wild-type and engrafted groups, deficient versus wild-type and engrafted groups individually, and engrafted versus wild-type groups. For comparisons versus the deficient response, enriched pathways corroborated with previously made differential gene expression associations to pro-inflammatory processes. Specifically, enrichment consistent in comparisons with the deficient samples notably highlighted upregulation of early or pro-inflammatory immune responses including inflammatory responses, neutrophil degranulation, innate immune system, IL-17 and TNF signaling pathways, and viral protein-like responses (Fig. 6a). Similarly, several independent pairwise comparisons against the deficient samples showed upregulation of pro-inflammatory TNF, and NF-kappa B and NOD-like receptor signaling pathways (Fig. 6b).

**Fig. 6:**
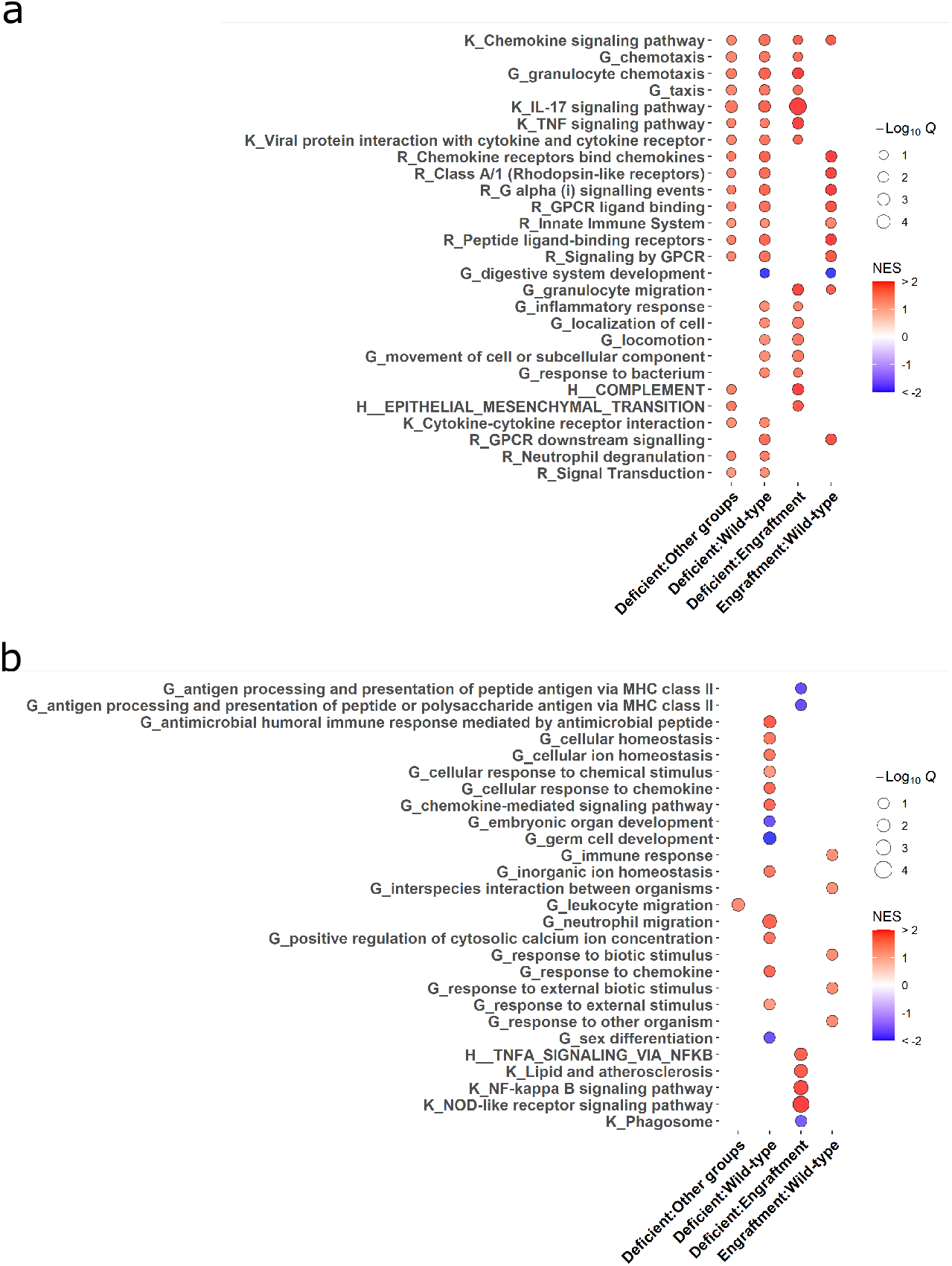
Mast cell-deficient male mice maintain late pro-inflammatory response to biomaterial implantation while mast cell engraftment restores physiological cellular responses to biomaterial stimuli. Bubble plot from gene set enrichment analysis for enriched pathways **(a)** consistent or **(b)** independently enriched amongst different group comparisons. Assessed databases included simplified biological process GO terms (“G_…”), KEGG pathways (“K_…”), Reactome pathways (“R_…”), and MSigDB hallmark gene sets (“H_…”). Color of each point represents magnitude and directionality of pathway based on normalized enrichment score (NES) and point size representing negative of the log_10_ transformed q-value. Results from n = 8 per group.

In contrast, enriched gene sets for the engrafted versus wild-type responses showed a lack of significant upregulation along the majority of these pro-inflammatory pathways except for potential upregulation associated with the innate immune system and granulocyte migration (Fig. 6a). Instead, enrichment consistent with the other comparisons mainly consisted of signaling and receptor binding pathways such as related to generalized chemokine and G protein signaling related pathways (Fig. 6a). Independent enrichment pathways for engrafted versus wild-type comparison also showed upregulation of general immune and responses associated to biotic stimuli (Fig. 6b), suggesting a more normalized immune response profile similar to the wild-type group. Ultimately, these results further supported the significant contribution of mast cell presence to the pro-inflammatory transition of the ECM biomaterial induced immune response.

## Discussion

Biomaterials have gained momentum as a pro-regenerative therapy through modulation of the innate and adaptive immune response. In particular, macrophage^7^ and T cell^11^ contributions have been a focus of studying the induced tissue remodeling progression. However, the immune response consists of a wide breadth of immune cell populations and discoveries of significant contributions of understudied cell populations to immunotherapies have highlighted the need to expand our investigative focus^45^. In this study, we focused our investigation into the contribution of mast cells. Increased numbers of tryptase positive mast cells had been identified following injection of an ECM hydrogel in a rat myocardial infarction model^27^; however, to the best of our knowledge, no previous studies have evaluated the contribution of mast cells to the immunomodulatory response of decellularized ECM biomaterials. Using a subcutaneous implant model, which can recapitulate the pro-remodeling immune cell polarization response of decellularized ECM materials seen in disease or injury models^8,29^ and allows for easy isolation of biomaterial scaffolds, we found non-degranulated mast cells in the nearby surrounding tissue, at the bioscaffold-tissue border, and in the core of the material, contrasting the degranulated states observed with synthetic biomaterial implants^3,16,46^.

Focusing on mast cell’s role in immune cell recruitment and influence by releasing immunoregulatory factors, implantation of the ECM biomaterial in male and female Kit^W-sh^ mast cell deficient mice demonstrated similar or greater total and immune subpopulation specific infiltration. This was observed despite the lack of mast cells releasing factors such as histamine for vasodilation and chemokines that are known to contribute for driving the early immune response, particularly for neutrophils^48,49^. Given that bioactive ECM components contain signals for promoting immune cell recruitment and migration^50^ or could potentially interact with other tissue resident immune cell populations, naturally derived ECM biomaterials do not appear dependent on mast cells for early immune cell recruitment. Furthermore, although some evidence of neutrophilia has been previously observed in this mast cell deficient model^51^, it did not manifest in significant differences in localized neutrophil infiltration in this experimental context.

Effects on temporal dynamics of macrophage polarization were observed by two independent methods that determined the ratio of alternatively activated M2 and classically activated M1 macrophages significantly differed between the wild-type and mast cell deficient mice. These differences were reversed by engraftment of *in vitro* differentiated bone marrow derived mast cells in deficient mice supporting differences were caused by mast cell presence. Some discrepancy in the timing of the observed M2 shifts was seen between the two assessments, which might occur from preferential expression amongst M2 subpopulations for certain M2 markers and stimuli specific expression patterns over time^52^.

Several differences also existed based on sex as male mice showed this dysregulation at the later day 11 timepoint while female mice showed altered ratios at the early day 3 timepoint. This drove additional investigation on the gene expression of cytokines and female hormone receptors, which determined increased expression of M1 marker Nos2 and estrogen receptor Esr2 in female deficient mice compared to wild-type. Though these results could be considered contradictory to flow cytometry results showing greater M2 profile in deficient mice at the day 3 timepoint, studies have demonstrated specific influence of female hormones on gene expression of Nos2^44,53^, specifically, with Esr2 increasing Nos2 expression in immune cells and other cell populations such as vascular smooth muscle cells^54,55^. This relationship could suggest that this classic M1 marker is not representative of the macrophage response in female mast cell deficient mice, which might be distinctly upregulated due to mast cells also being influenced by sex specific hormones^56^. Although outside the scope of this study to fully delineate the differences in biomaterial response based on sex, it is well known that immune response characteristics and tissue disease prevalence, pathology, and symptoms differ based on sex^1,57^. Thus, these results highlight, especially when considering future translation of these biomaterial therapeutics and variability at the clinical stage, the need for further investigation into the understudied field of sex/hormone specific responses to biomaterial therapeutics.

For male mice samples, differences in pro-inflammatory gene expression profiles in the mast cell deficient mice compared to both wild-type and mast cell engrafted mice corroborated the altered polarization determined from macrophage assessments. This was especially distinct between the mast cell deficient and engrafted mice considering the specific restoration of localized mast cell presence led to a limited set of differentially regulated genes that were predominately pro-inflammatory markers: chemokines (Ccl2, Ccl3, Ccl7, Cxcl1, Cxcl3), cytokines (Il1b, Il1rl1, Il6, Il33, Tnf), alarmins (S100a8, S100a9), and regulatory aids in pro-inflammatory propagation (Ptgs2, Sele, Trem1). This was also highlighted in the enrichment analyses where deficient samples were upregulated in early or pro-inflammatory related gene sets amongst the various group comparisons. Ultimately, these comparisons demonstrated that mast cell presence aids in the temporal dynamics and regulation of the pro-inflammatory to pro-remodeling response for an ECM biomaterial.

Considering the consistency of pro-inflammatory markers differing in the deficient male mouse response compared to either wild-type and engrafted responses, comparisons were also made between wild-type and engrafted groups to evaluate the degree mast cell engraftment restored the wild-type response. These results also provided evidence of whether specific localization of mast cells around the biomaterial space were beneficial or detrimental. Based on differential analysis, significant genes were mainly associated with immune cell regulation, recruitment, differentiation, and activation along with general cell metabolic and signaling genes while lacking enrichment of distinct pro-inflammatory markers. Thus, while some differential genes and pathways were present between the wild-type and engrafted samples, the pro-inflammatory response was normalized to the wild-type response with mast cell engraftment. Several differentially regulated genes that were not normalized with mast cell engraftment, such as Tslp and Il33, potentially demonstrate maintained upregulation of beneficial regulatory responses as these genes have been specific therapeutic targets in immunomodulatory biomaterial therapy studies for regulating chronic inflammatory responses^58–60^. These results go in line with the other analyses showing increased pro-remodeling type responses with localized mast cell delivery such as an increased CD206^+^ macrophage presence in engrafted compared to the wild-type response. Thus, localized mast cell presence can provide contributions to the induced immune response extending beyond just mitigating pro-inflammatory responses by promoting beneficial regulatory responses in the biomaterial immune microenvironment.

One limitation of this study is that mast cell deficiency inherently influences tissue remodeling and function with injury^61,62^, thus, making it difficult to definitively determine whether mast cell deficiency influences biomaterial induced endogenous tissue repair responses when inherent markers of chronic tissue diseases such as function or fibrosis are significantly affected. With more specific understanding of mast cell characteristics *in vivo*, genetic manipulations to specifically manipulate mast cell phenotype could be performed such as crossing *Mcpt5*-*Cre* transgenic mice with *IL-10*^fl/fl^ for mast cell specific knockout of IL10^63^ and other similar strategies^64^. However, it is known that some off-targeting effects have been observed with these mouse models^64^, thus, this is still an actively developing field to improve mast cell specific manipulations in animal models.

Overall, in contrast to previous *in vivo* studies of the mast cell response to biomaterials, which showed involvement in acute inflammation and fibrous capsule formation^3,16^, this study demonstrated that mast cells contribute to the dynamic immunomodulatory response to pro-regenerative biomaterial scaffolds. This work highlights the need to examine the role of other immune cells beyond macrophages and T cells as well as sex differences in pro-regenerative therapies, and supports the potential of mast cells as a critical immune regulatory element for stimulating endogenous tissue repair.

## Materials and Methods

All procedures in this study were performed in accordance with the guidelines established by the Committee on Animal Research at the University of California, San Diego and the Association for the Assessment and Accreditation of Laboratory Animal Care.

### ECM Hydrogel Preparation

An ECM hydrogel was generated from porcine left ventricular tissue based on previously established protocols^65^. In brief, porcine left ventricular tissue was isolated and minced into small pieces. The tissue was decellularized under mechanical agitation in a solution of phosphate buffered saline (PBS) containing 1% (wt/vol) sodium dodecyl sulfate (SDS) (Fischer Scientific, Fair Lawn, NJ) with 0.5% 10,000 U/mL penicillin streptomycin (PS) (Gibco, Life Technologies, Grand Island, NY) until fully decellularized based on previously established criteria^65^. Once decellularized, the decellularized ECM was thoroughly rinsed to remove residual SDS, lyophilized, and milled into a fine powder. ECM powder was partially digested with 1 mg/mL pepsin in 0.1M HCL solution for 48 hours before solution was neutralized to pH of 7.4 and reconstituted to physiological salt concentrations. Partially digested ECM solution was aliquoted, lyophilized and stored at −80**°**C until re-suspending with sterile water prior to injection which self-assemble into hydrogels *in vivo*. Non-decellularized material was created as previously described^8^, where a one day rinse in PBS with PS replaced the multiple days of decellularization with SDS. The non-decellularized material was then similarly processed into an injectable form as the decellularized material described above.

### Mast Cell-Deficient Mouse Model

Homozygous mast cell-deficient mouse, w-sh mice: B6.Cg-*Kit*^*W*-*sh*^/HNihrJaeBsmJ (Jackson Laboratory, Sacramento, CA) were breed in UCSD vivarium up to 11-13 weeks before being designated for subcutaneous procedures. Both male and female mice were utilized for investigation. Age matched wild-type C57BL6/J mice were used as control animals.

### Bone Marrow Derived Differentiated Mast Cell Culture

Bone marrow derived cells were harvested from wild-type C57BL6/J mice that were euthanized by CO2 and cervical dislocation. Each femur was isolated and bone marrow was flushed with 10 mL of media, collected and pipetted into culture flasks incubated in cell culture incubator at 37°C and 5% CO_2_. Mast cell differentiation media consisting of DMEM media (Gibco) with 10% FBS (Gibco), 1% penicillin streptomycin (Gibco), sterile filtered conditioned culture medium from D11 hybridoma cells as a source of IL3, 100 uM MEM nonessential amino acids (Lonza), 1mM sodium pyruvate (Gibco), 0.1 mM HEPES (Gibco), and 55 uM β-mercaptoethanol (Gibco). Non-adherent cells were passaged at day 3, 7, 10, 14, 18, 22 and 28 before differentiation of mast cells was assessed by flow cytometry with antibody markers for CD117 APC (2B8, Biolegend, San Diego, CA) and FcεRI PE (MAR-1, Biolegend). Cells at greater than ∼95% purity were utilized up to 4-6 weeks from initial isolation before mast cell functionality was expected to subside.

For assessment of cell viability, 100,000 mast cells were plated into a 48 well plate (n = 4) with ECM hydrogel material at nongelling concentrations of 0.6 mg/mL or neutralized Collagen I, Rat Tail at concentrations of equivalent stiffness and collagen content of 0.25 mg/mL concentration^66^ (Corning®, Corning, NY) doped into culture media. Media only culture was used as a control. AlamarBlue™ Cell Viability Reagent (Invitrogen, Waltham, MA) was added to each well at a tenth the volume totaling 400 μL solution, and cells were incubated in a 37°C, 5% CO_2_ cell culture incubator. At 0, 2, 4, and 8 hours after plate set-up, fluorescent plate readings for alamarBlue signal at 560 nm excitation and 590 emission were read on a Synergy™ H4 multi-mode microplate reader and Gen5™ software (Biotek®. Wilnooski, VT). Experimental set-up was repeated to demonstrated relative reproducibility of the results.

### Mast Cell-Engrafted Model Generation

A mast cell-engrafted model was generated using a previously described protocol^26,42^. For injection procedures, animals were briefly put under anesthesia at 2% isoflurane on a nose cone and mice went placed upright to expose the dorsal region. The dorsal region was shaved and cleaned with 3 intervals of betadine and 70% ethanol. Mast cell-engrafted mice were created by delivering eight to ten 100 μL subcutaneous injections of bone marrow differentiated mast cells in DMEM totaling 4 million cells. Injections were at evenly spaced intervals throughout the dorsal region of 4-6 week old deficient mice. Subsequent procedures were done in engrafted mice at 11-13 weeks old. Toluidine blue staining of subcutaneous tissue and injections confirmed presence of mast cells in engrafted mice at the time of subcutaneous procedures and harvest (Supplementary Fig. 6). For comparisons with engrafted mice, age matched mast cell deficient mice, injected with DMEM alone at similar timepoints for comparative studies, and wild-type mice were utilized as controls.

### Biomaterial Injection and Harvesting

For biomaterial injections, animals were briefly put under anesthesia using 2% isoflurane on a nose cone and each mouse was injected with ECM hydrogel material receiving two evenly spaced 200 μL subcutaneous injections in the upper and lower dorsal region. At one, three, seven, and eleven days post-injection, mice were euthanized by CO_2_ and cervical dislocation. Timepoints were selected based on expectations of the immune response being at the pro-inflammatory and pro-remodeling phases based on previous studies^8^ along with observation of consistent material retention and clear visibility up to the day eleven timepoint. General dorsal skin tissue was cut and flipped to expose the underside to observe location and area of material injection. The injections, along with neighboring dermal tissue, were excised. Each harvested injections were divided into multiple parts for analysis by flow cytometry, staining or gene expression analysis as described below (n= 3-8 mice per group, 6-16 injections per group). For timepoints with notable differences between the deficient and wild-type responses, samples were collected across 2-3 experimental batches of animals for demonstrating consistency of results.

### Flow Cytometry

Pieces from multiple excised subcutaneous injections per animal were pooled and minced in ice-cold HBSS (Gibco) and enzymatically digested in a solution consisting of 1:1 solution of HBSS (calcium and magnesium supplemented) and 1% bovine serum albumin in PBS with 1 μM HEPES (Gibco), 300 U/mL collagenase type IV (Worthington Biochemical), 60 U/mL hyaluronidase (Sigma-Aldrich) and 10 U/mL DNase I (Sigma-Aldrich). Material in enzymatic digestion solution were incubated at 37°C under mechanical agitation at 750 rpm on a thermomixer (Benchmark Scientific) for 45 minutes. Solutions were then kept in ice and FACs buffer consisting of 1% bovine serum albumin and 1mM EDTA in DPBS lacking calcium and magnesium added to inhibit further enzyme reaction. Digested tissue was filtered through a 100μm cell strainer. Cells were centrifuged at 400 rcf centrifugation at 4°C and resuspended in HBSS. Cell suspension was stained with LIVE/DEAD™ Fixable Aqua (ThermoFisher Scientific) for 10 minutes at 4°C and excess dye was quenched with FACs buffer. Cells were fixed and permeabilized by BD Cytofix/Cytoperm™ Buffer (BD Biosciences) for 10 minutes and washed in BD Perm/Wash™ Buffer (BD Biosciences). Cells were counted by hemocytometer and stained with antibody panels for immune cell subpopulations. General immune cell populations were stained with the following antibody panel: CD11c BV421 (N418, Biolegend), F4/80 BUV395 (T45-2342, BD Biosciences), CD3 PerCp/Cy5.5 (17A2, Biolegend), CD117 APC (2B8, Biolegend), FcεRI PE (MAR-1, Biolegend), and CD19 APC-Cy7 (6D5, Biolegend). An antibody panel for macrophage and T cell polarization was also stained for consisting of: F4/80 BUV395 (T45-2342, BD Biosciences), CD86 BV786 (GL-1, Biolegend), CD206 PE (C068C2, Biolegend), CD3 PerCp/Cy5.5 (17A2, Biolegend), CD4 APC (GK1.5, Biolegend), CD8a Alexa Fluor 488 (53-6.7, Biolegend), and FoxP3 BV421 (MF-14, Biolegend). Antibodies and IgG isotype controls were stained in BD Perm/Wash™ Buffer for 30 minutes at 4°C before rinsing and resuspending in FACs buffer consisting of 1% bovine serum albumin and 1 mM EDTA in PBS for analysis. Stained cells were analyzed on a BD FACSCanto™ II and BD LSRFortessa™ X-20 (BD Biosciences). Gating was set based on positive, isotype and fluorescence minus one controls utilizing a mixed single cell suspension control sample derived from seven day post-injection biomaterial, isolated spleen cells, and four week differentiated mast cells to ensure all cells of interest were present in distinguishable amounts. Gating and flow data were processed in FlowJo v10.6.2 (FlowJo LLC, Ashland, OR).

### Quantitative Real-Time Polymerase Chain Reaction (qRT-PCR)

Each flash frozen or RNAlater™ (Invitrogen) treated subcutaneous injection sample was homogenized with a mechanical rotator and then run through a RNeasy Mini kit (Qiagen, Germantown, MD) to extract RNA based on manufacturer instructions with an on-column DNase I digestion (Qiagen) for minimizing genomic DNA contamination. Superscript IV Reverse Transcriptase kit (Applied Biosystems, Foster City, MA) was used to synthesize cDNA with thermocycler settings of 65°C for 5 minutes, 23°C for 10 minutes, 55°C for 10 minutes and 80°C for 10 minutes. Eva Green Master Mix (Biotium, Fremont, CA) was used with custom-made forward and reverse primers (Table S1) at a final concentration of 0.2 μM for qPCR reactions. Samples were run in technical duplicate along with negative controls without template cDNA to confirm lack of contamination from qPCR reagents. PCR reactions were run on a CFX95™ Real-Time System (Bio-Rad, Hercules, CA) with the following thermal cycler settings: 30s at 50°C, 2 min at 95°C, 40 cycles of 10s at 95°C, and 30s at 55-65°C based on pre-determined optimal primer efficiency amplification temperature. After completing 40 cycles of PCR amplification, automated melting curve analysis, consisting of increasing the thermal cycler temperature from 50**°**C to 95**°**C at 5**°**C increments lasting 5s each, for confirming singular amplicon product in each reaction vessel. Bio-Rad CFX Manager™ 3.0 (Bio-Rad) was used for determining cycle threshold values from recorded signal based on a preset threshold.

### Nanostring Multiplex Gene Expression Analysis

For comprehensive evaluation of the whole immune profile, equal amounts of RNA sample across subcutaneous injections per animal were pooled (n = 8 pooled subcutaneous injections representing mouse replicates per group) and were analyzed by Nanostring nCounter® MAX Analysis System with nCounter® Immunology Panel (Mouse) allowing for multiplexed assessment of 546 genes^67,68^. Samples were processed according to manufacturer instructions. In brief, RNA sample concentrations were measured on a Qubit 3.0 Fluorometer with a Qubit™ RNA HS Assay kit. 70 μL of hybridization buffer was mixed with Immunology Panel Reporter CodeSet solution, and 8 μL of this master mix was mixed in a separate reaction vessel with 100 ng of RNA per tissue sample and RNA-free water up to 13 μL total. 2 μL of Capture ProbeSet was added to each vessel, mixed and placed on a thermocycler at 65°C for 16-48 hours before being maintained at 4°C for less than 24 hours. Nanostring nCounter Prep Station performed automated fluidic sample processing to purify and immobilize hybridized sample to cartridge surface. Digital barcode reads were analyzed by Nanostring nCounter® Digital Analyzer.

Results were analyzed by manufacturer nSolver™ Analysis Software 4.0 and custom R scripts under R versions 4.04. Gene expression normalization and differential expression was determined by the NanostringDiff package^69^ with genes with average probe count one standard deviation below average negative control probe count excluded from analysis. Significance was set with a false discovery rate Benjamini-Hochberg method correction for calculating significance based on q-value < 0.05 and an absolute log2 fold change ≥ 1. Heatmap of significant differentially expressed genes was displayed with the pheatmap package^70^. Volcano plots created with the EnhancedVolcano package^71^ displaying the top 25 genes based on aggregated ranking for lowest q-value and greatest absolute log2FC using the RankAggreg package^72^. Gene set enrichment analysis was performed with the clusterprofiler package^73^ across the following databases: Gene Ontology (GO) terms^74,75^ with redundancy reduced with the GOSemSim package^76^, Kyoto Encyclopedia Encyclopedia of Genes and Genomes (KEGG) pathways^77^, Reactome pathways^78^ with the ReactomePA package^79^, and hallmark gene sets from the Broad Institute and UCSD derived MSigDB^80^ with the msigdbr package^81^. KEGG pathways related to disease gene sets were excluded with the gageData package^82^, at least three genes were required to be enriched for gene sets, and significance cut-off for enrichment analysis was set at q-value < 0.05 with a false discovery rate by Benjamini-Hochberg correction.

### Immunohistochemistry

Harvested samples were embedded into Tissue-Tek O.C.T. Compound (Sakura®, Torrance, CA) for cryosectioning. Cryosections from two-three different evenly spaced locations were used for all immunohistochemistry, and injections too small to obtain at least two locations (around less than 300 μm total width) were not processed. Transverse sections were taken to obtain a cross-section of the biomaterial injection and neighboring dermal tissue. Histological evaluation of total cell and mast cell infiltration were taken from sections stained with hematoxylin and eosin (H&E) or Toluidine Blue Stain, 1% w/v (Ricca Chemical, Arlington, TX) and scanned with a Leica Aperio ScanScope® CS^2^ (Leica, Buffalo Brove, IL) system. For fluorescent staining, slides were fixed with 4% paraformaldehyde in PBS (Thermo Scientific™) and blocked with a buffered solution containing 3% bovine serum albumin (Gemini Bio-Products, Inc., West Sacramento, CA) and 0.1% Triton X-100 (Sigma) in PBS. The primary antibodies, pan-macrophage marker anti-mouse F4/80 (BM8, 14-4801-82, eBioscience, San Diego, CA) and anti-mouse Ly-6G/Ly-6C (RB6-8C5, 14-5931-82, eBioscience), were incubated for 12-18 hours at 4°C: Secondary antibody Alexa Fluor 488 with Hoechst 33342 (Thermo Scientific™) counterstain was incubated for 30-40 minutes at room temperature. Coverslips were mounted with Fluoromount™ Aqueous Mounting Medium (Sigma) and allowed to dry protected from light. Fluorescent images were scanned with the Leica Ariol® DM6000B system (Leica). Cellular density and co-staining quantification were done with custom MATLAB scripts (Mathworks, Natick, MA). Nuclei quantification was averaged across at least two locations per injection for nuclei staining (n = 8-10 injection replicates or 4-5 mouse replicates per group) while locations across both injections were averaged for co-staining analysis due to greater variability and background signal from material non-specific staining (n = 4-5 mice replicates per group).

### Statistical Analysis

All data and plots are presented as mean ± SD unless noted otherwise. Statistics were performed in Prism 8 (GraphPad Software, San Diego, CA) or custom R scripts with publicly available packages. For two group assessments, an unpaired student’s *t*-test was performed while assessments across three or more groups was evaluated with a one-way ANOVA with a post-hoc Tukey with significance taken at *p* < 0.05. For analysis of differential expression of Nanostring data, a false discovery rate threshold of *q* < 0.05 based on the Benjamini-Hochberg method and an absolute log2 fold change threshold of greater than or equal to 1 were utilized. Gene set enrichment analysis utilized a p-value < 0.05.

## Data and materials availability

Raw Nanostring Mouse Immunology Panel counts, genes analyzed post-filtration, differential expression analyses, and gene set enrichment analysis determined for conclusions in the paper are available in the Supplementary files. Additional data may be available from the corresponding author upon reasonable request.

## Code availability

Custom scripts utilized for data analysis utilized publicly available packages as described in the manuscript, and may be available from the corresponding author upon reasonable request.

## Acknowledgements

The authors would like to thank Drs. Thomas Gilbert and Adam Young from ACell, Inc. for kindly providing the milled urinary bladder matrix. We would also like to acknowledge Dr. Elsa Molina, Director of the UC San Diego Stem Cell Genomics Core, for technical assistance of NanoString experiments, and Jesus Olvera, Cody Fine, and Vu Nguyen of the UC San Diego Human Embryonic Stem Cell Core Facility for technical assistance of flow cytometry experiments with all made possible in part by the CIRM Major Facilities grant (FA1-00607) to the Sanford Consortium for Regenerative Medicine.

## Funding

This work was supported by the NIH NHLBI (R01HL113468, R01HL146147). R.M.W. and J.L.U. were supported by NIH NHLBI pre-doctoral fellowships (F31HL137347, F31HL136082). J.M.M. was funded by the NIH NHLBI as a T32 training grant recipient (T32HL105373).

## Author contributions

R.M.W contributed to the project conceptualization and design, experimental work, data collection and analysis, and manuscript write-up. J.M.M contributed to project design, experimental work, data collection, and manuscript feedback. J.L.U. contributed to project conceptualization and design, and experimental work. Yu K. contributed to initial training and feedback on mast cell methods. Yuko K. contributed to initial training, reagents for mast cell work, and feedback on mast cell methods. T.K. contributed guidance for project design and experimental work, feedback on data, and evaluation of manuscript. K.L.C. contributed to project conceptualization, guidance for project design and experimental work, feedback on data, and evaluation of manuscript. All authors edited the manuscript.

## Competing interests

KLC is co-founder, consultant, board member, and holds equity interest in Ventrix, Inc. Other authors declare that they have no competing interests.

