## Supplemental Figures for "Immunomodulatory Contribution of Mast Cells to the Regenerative Biomaterial Microenvironment"

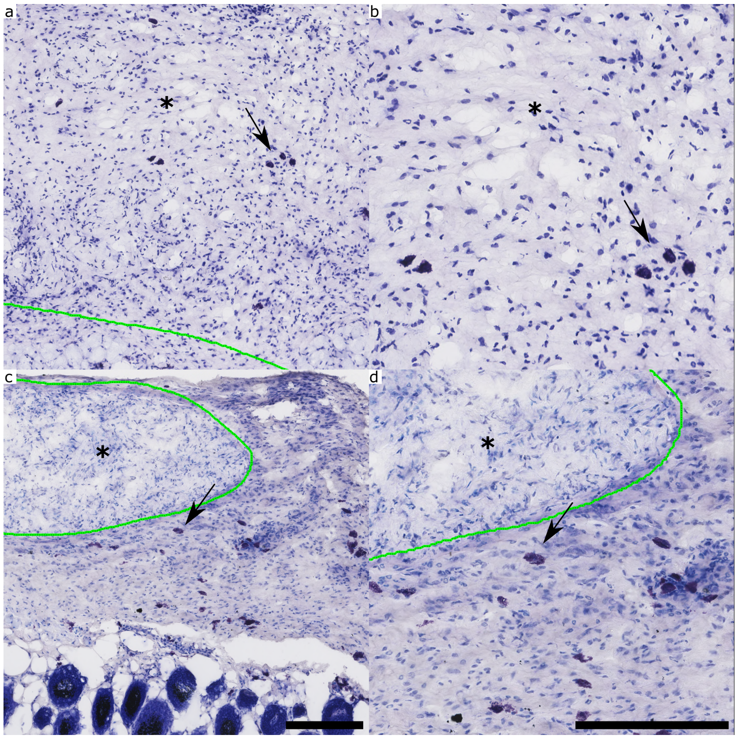


**Supplementary Figure 1: Toluidine blue staining of subcutaneously injected ECM hydrogel and urinary bladder matrix powder.** Representative and zoomed images of toluidine blue staining of **(a, b)** ECM hydrogel scaffold and **(c, d)** milled urinary bladder matrix (material outlined in green with asterisk on side where material is present) and neighboring dermal tissue in male wild-type 3 days post-injection. Mast cells staining is indicated by a black arrow. Scale bar is 200 μm.

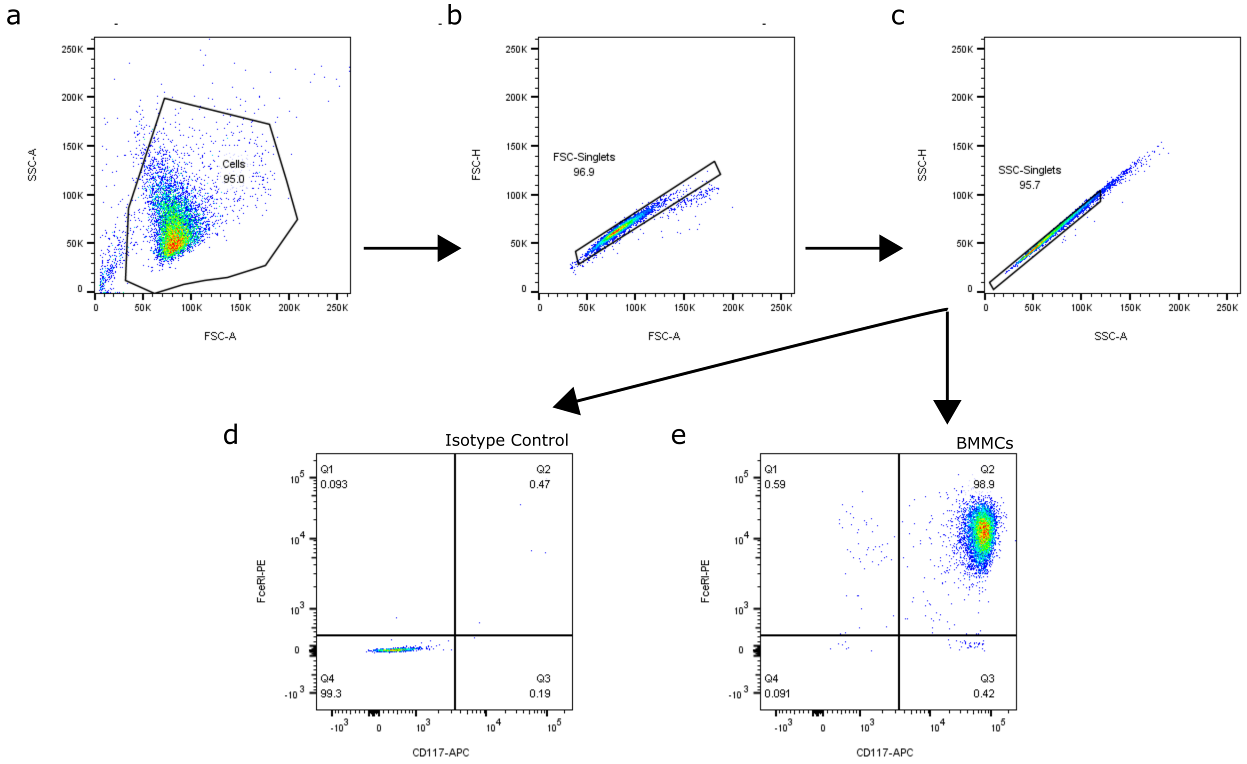


**Supplementary Figure 2: Gating scheme for bone marrow derived differentiated mast cell culture purity.** Representative gating of mast cells for assessing purity at four to six weeks of differentiation protocol. (**a)** Cells and **(b-c)** double discrimination of singlets based on forward and side scatter. Quadrant gating cut-offs were set relative to **(d)** isotype control for **(e)** $CD117^{+}FcƐRI^{+}$ mast cells.

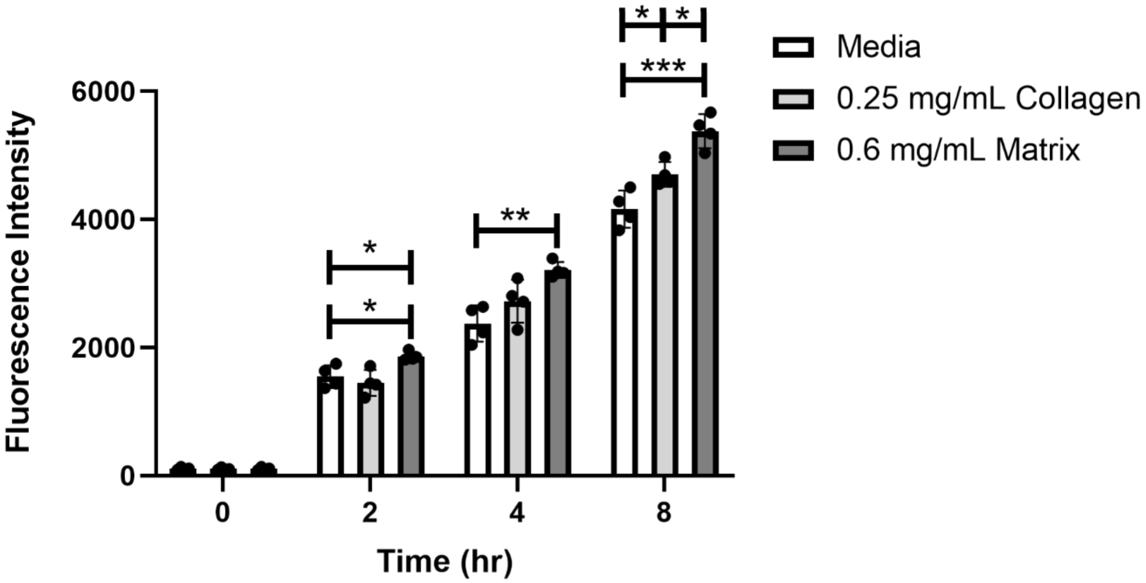


**Supplementary Figure 3: Viability assay of mast cells incubated with ECM hydrogel.** Four week differentiated bone marrow derived mast cells were incubated with ECM hydrogel material, collagen or culture media control. Collagen concentration was chosen based on previous studies showing analogous material properties to ECM hydrogels at this relative concentration[1]. AlamarBlue™ fluorescent readings were taken at 0, 2, 4, and 8 hours following plate set-up (n = 4 per group). (* $p < 0.05$, ** $p < 0.01$, *** $p < 0.001$).

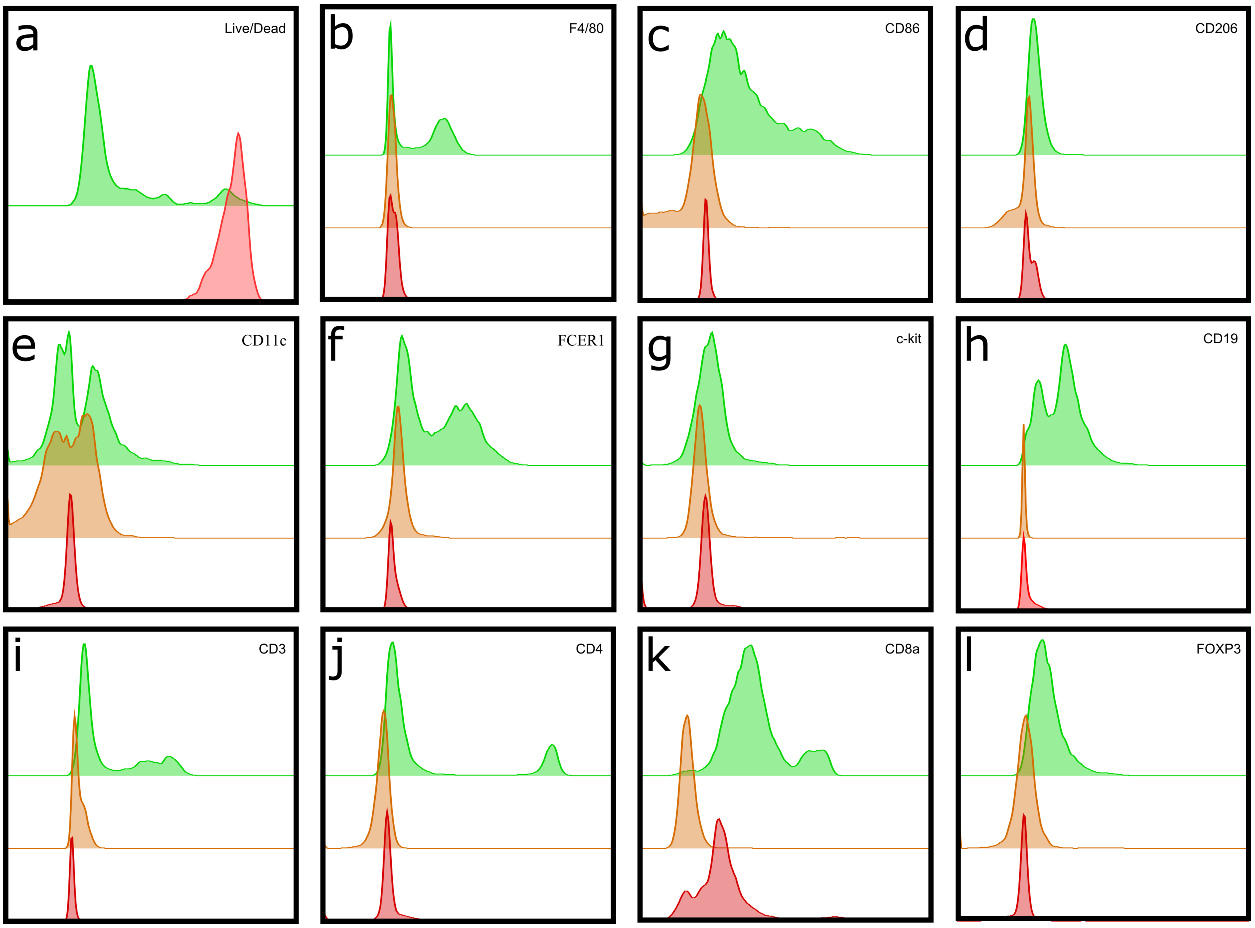


**Supplementary Figure 4: Gating controls for flow cytometry markers. (a)** LIVE, DEAD$^{TM}$ Aqua thresholds utilized ethanol fixed sample as control (red) for determining live cell gating (green). **(b-l)** Gating for fluorescent antibody markers was set for positive signal (green) based on isotype (red) and fluorescence minus one controls (orange).

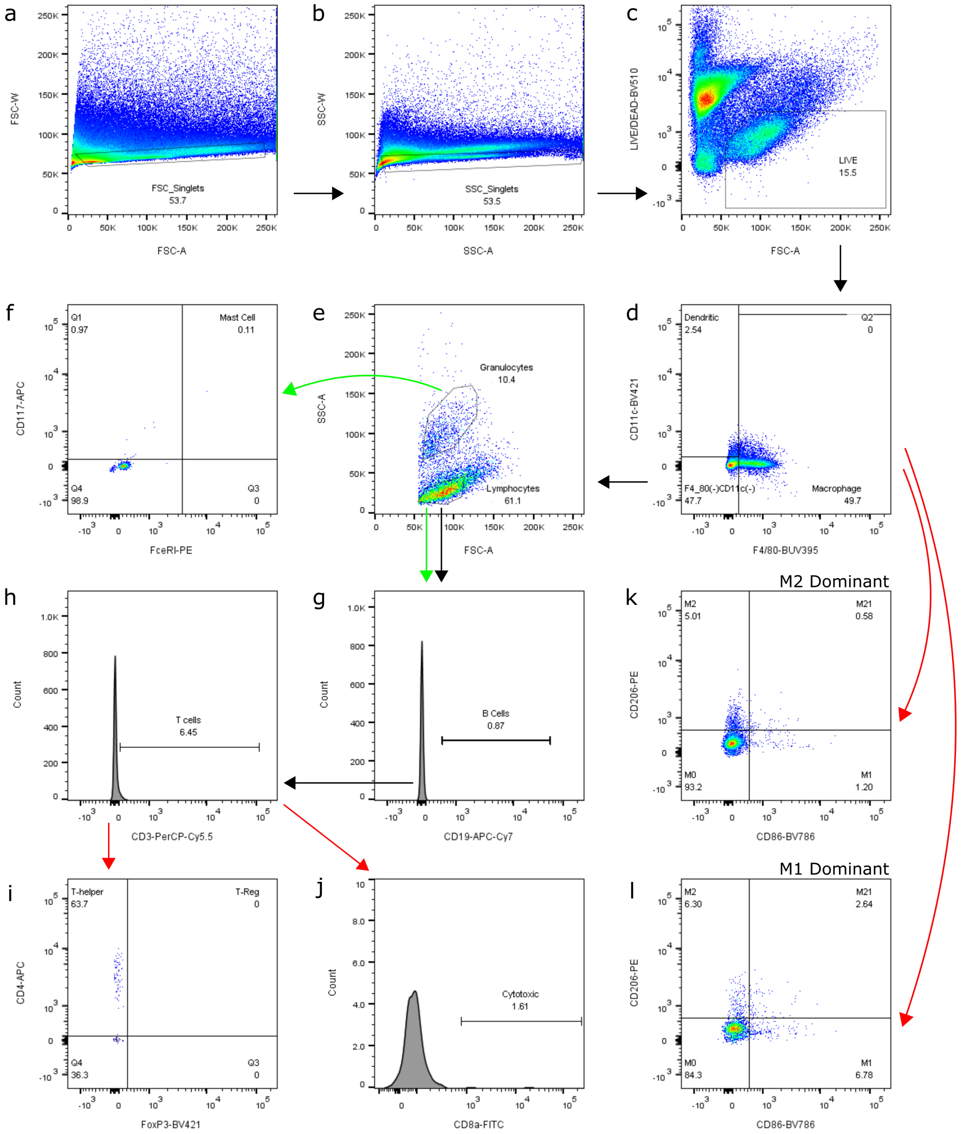


**Supplementary Figure 5: Gating Scheme for flow cytometry analysis.** Representative gating of single cell suspension isolated from subcutaneously injected ECM hydrogel. General gating for both panels (black arrow) started with double discrimination of singlets based on **(a)** forward and **(b)** side scatter. **(c)** Live cells were gated based on clustered from lack of stain uptake and forward scatter. **(d)** Macrophage and dendritic cell populations were gated based on F4/80$^+$ and CD11c$^+$, respectively. **(e)** F4/80$^-$CD11c$^-$ populations were separated into granulocyte and lymphocytes. For a general immune cell population panel (green arrow), granulocytes were assessed as **(f)** CD117$^+$FcƐRI$^+$ mast cells and lymphocytes for **(g)** CD19$^+$ B cells and **(h)** CD3$^+$CD19$^-$ for T cells. For immune cell polarization panel (red arrow), CD3+ T cells were assessed as **(i)** CD4$^+$FOXP3$^-$ T helper cells, CD4$^+$FOXP3$^+$ regulatory T cells, and **(j)** CD4$^-$FOXP3$^-$CD8a$^+$ cytotoxic T cells. F4/80$^+$ macrophage polarization were assessed for numbers of M1 CD86$^+$CD206$^-$ and M2 CD86$^-$CD206$^+$ markers with representative samples showing a **(k)** M2 dominant response versus a **(l)** M1 dominant response.

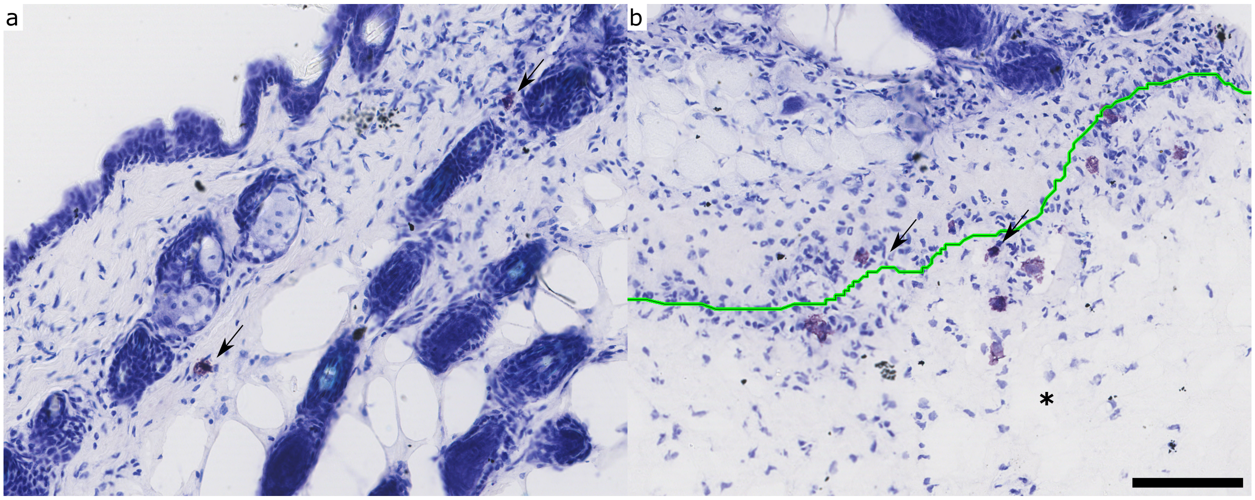


**Supplementary Figure 6: Mast cells engraftment in deficient mice.** Representative toluidine blue staining in 11-13 week old deficient mice following dorsal subcutaneous injection of *in vitro* differentiated bone marrow derived mast cells at 4-6 weeks old showing reconstitution of mast cells in the **(a)** skin and infiltrating into subcutaneously injected **(b)** ECM scaffold at 3 days post-injection (material outlined in green with asterisk on side where material is present). Mast cells staining is indicated by a black arrow with both resting and degranulated mast cells observed at this timepoint. Scale bar is 100 µm.

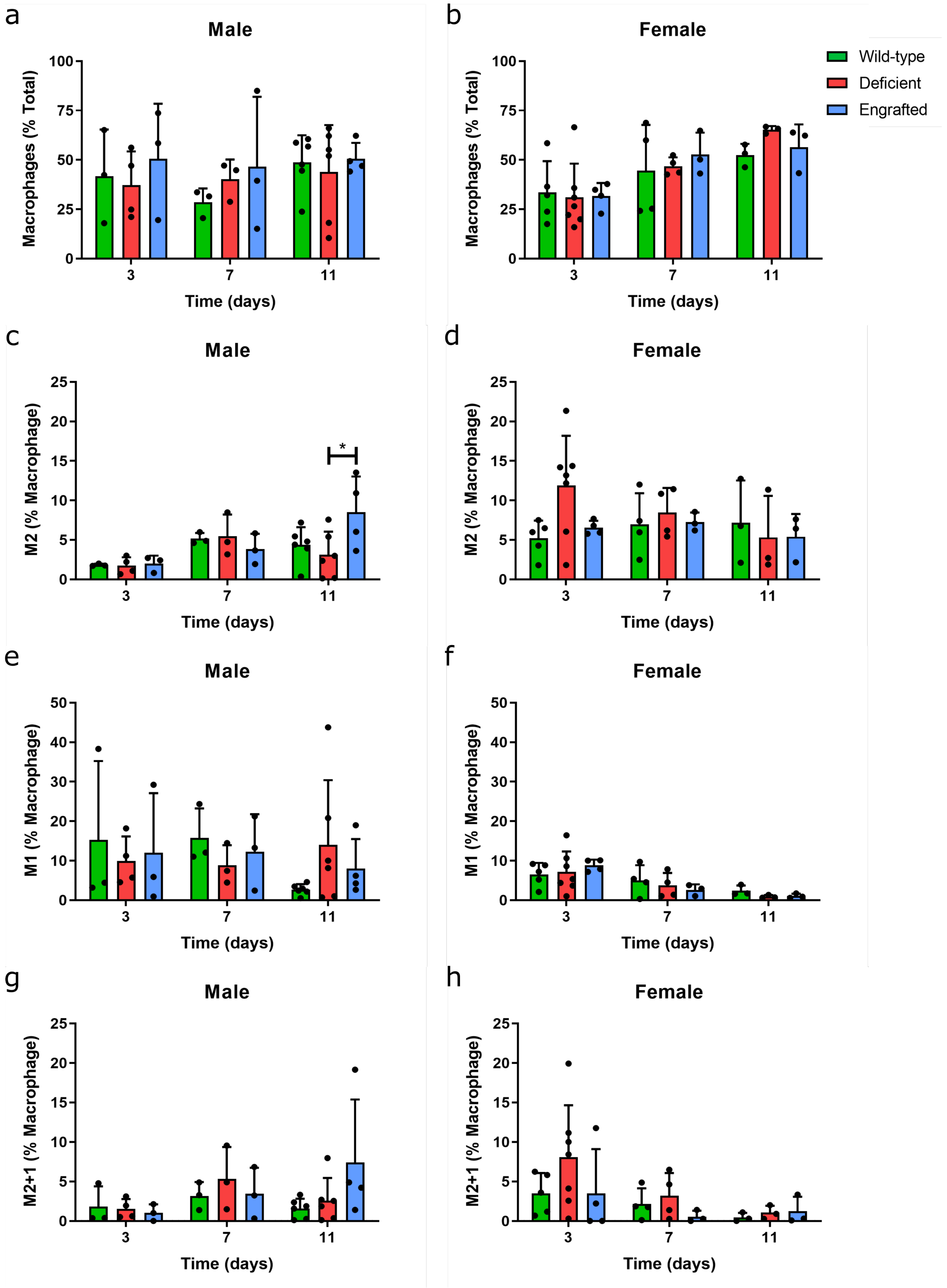


**Supplementary Figure 7: Flow cytometry of total and macrophage polarized subpopulations.** Percentages of total F4, 80⁺ macrophages relative to live cells **(a, b)**, and CD206⁺ M2 **(c, d)**, CD86⁺ M1 **(e, f)** and co-expressing CD206⁺CD86⁺ M2+1 **(g, h)** macrophages relative to total macrophages at three, seven, and eleven days post-injection between wild-type (green), deficient (red), and engrafted (blue) male **(a, c, e, g)** and female mice **(b, d, f, h)**. n = 3-6 per group.

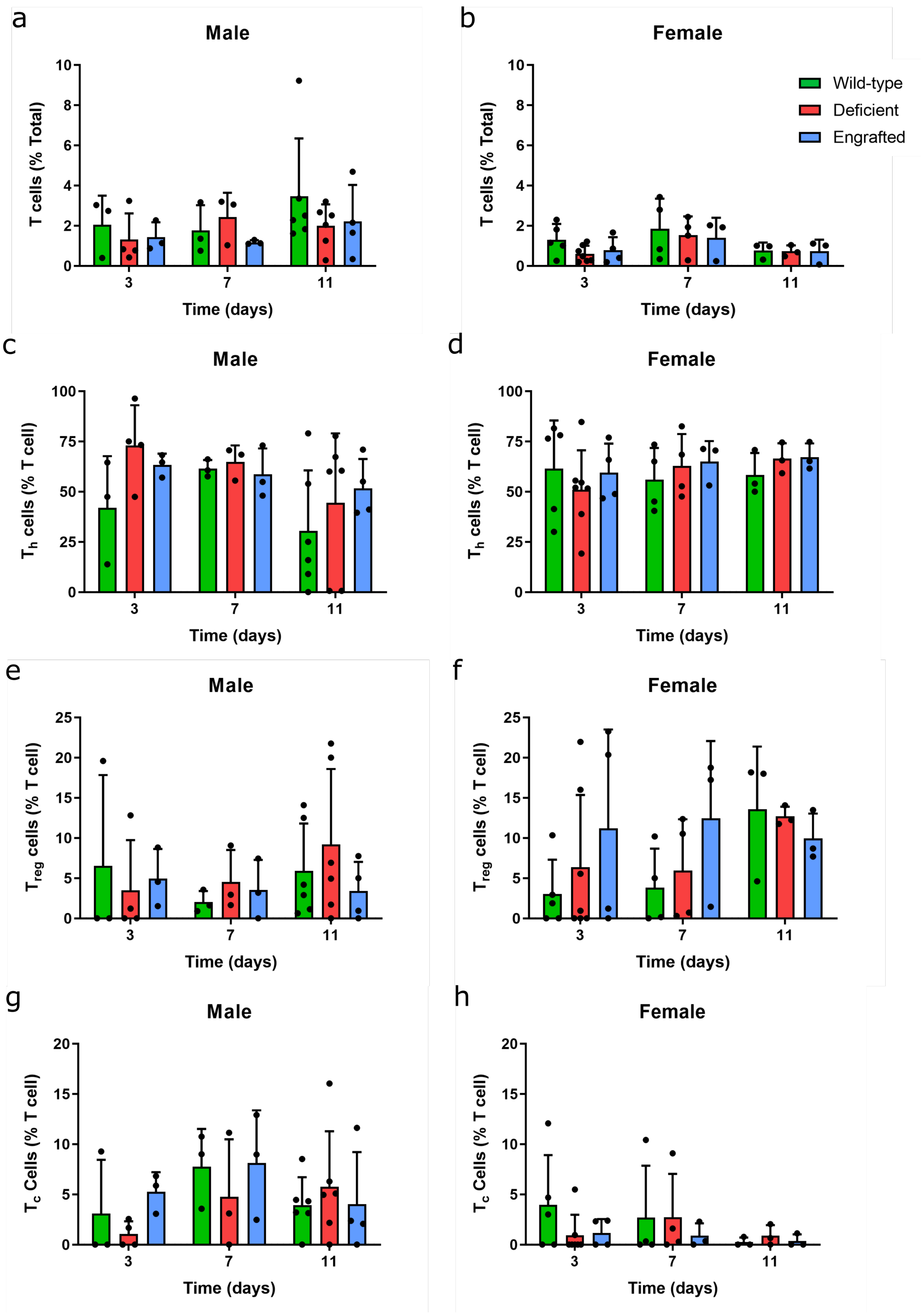


**Supplementary Figure 8: Flow cytometry of total T cells and differentiated subpopulations.** Percentages of total $CD3^+$ T cells relative to total live cells **(a, b)**, and $CD4^+$ T-helper ($T_h$) cells **(c, d)**, $CD4^+FOXP3^+$ regulatory T ($T_{reg}$) cells **(e, f)** and $CD8^+$ cytotoxic T ($T_c$) cells **(g, h)** relative to total T cells at three, seven, and eleven days post-injection between wild-type (green), deficient (red), and engrafted (blue) male **(a, c, e, g)** and female mice **(b, d, f, h)**. n = 3-6 per group.

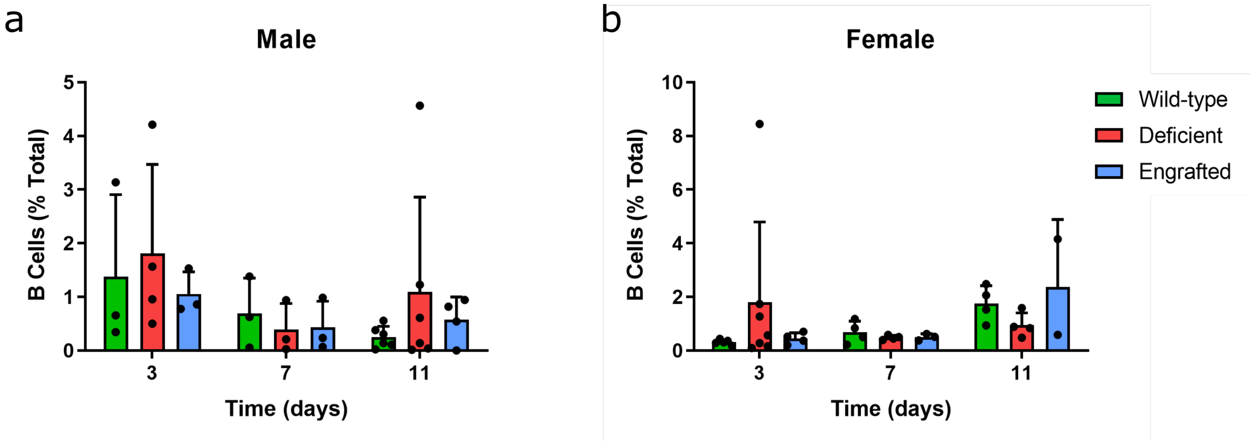


**Supplementary Figure 9: Flow cytometry of B-cells.** Percentages of $CD19^{+}$ B cells relative to total live cells at three, seven, and eleven days post-injection between wild-type (green), deficient (red), and engrafted (blue) male **(a)** and female mice **(b)**. n = 3-6 per group.

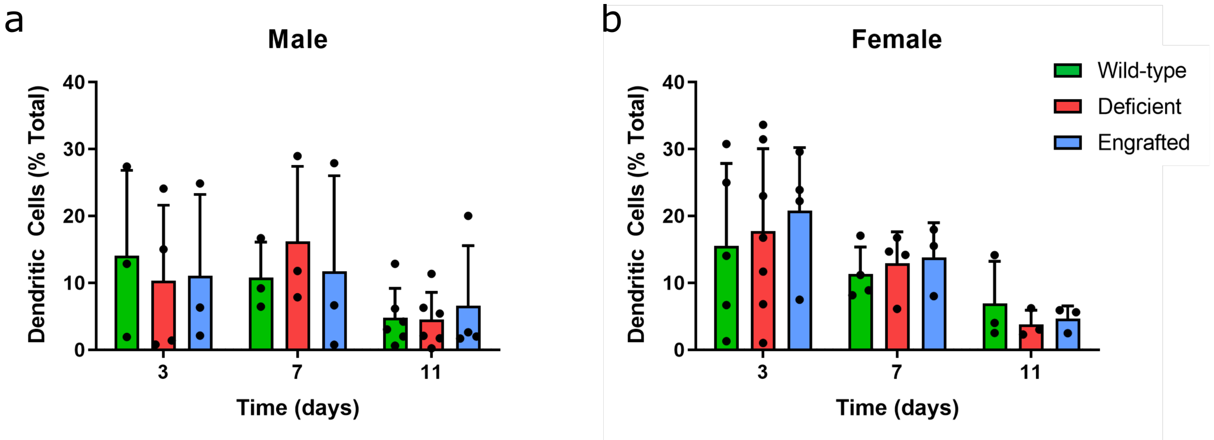


**Supplementary Figure 10: Flow cytometry of dendritic cells.** Percentages of CD11c$^{+}$F4, 80$^{-}$ dendritic cells relative to total live cells at three, seven, and eleven days post-injection between wild-type (green), deficient (red), and engrafted (blue) male **(a)** and female mice **(b)**. n = 3-6 per group.

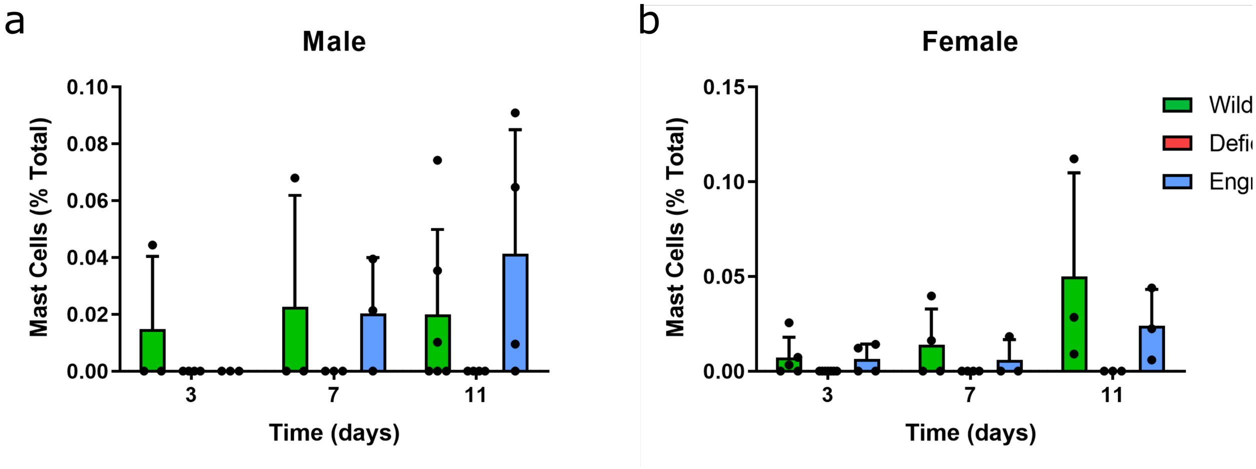


**Supplementary Figure 11: Flow cytometry of mast cells.** Percentages of CD117$^+$FcƐRI$^+$ mast cells relative to total live cells at three, seven, and eleven days post-injection between wild-type (green), deficient (red, not present), and engrafted (blue) male **(a)** and female mice **(b)**. n = 3-6 per group.

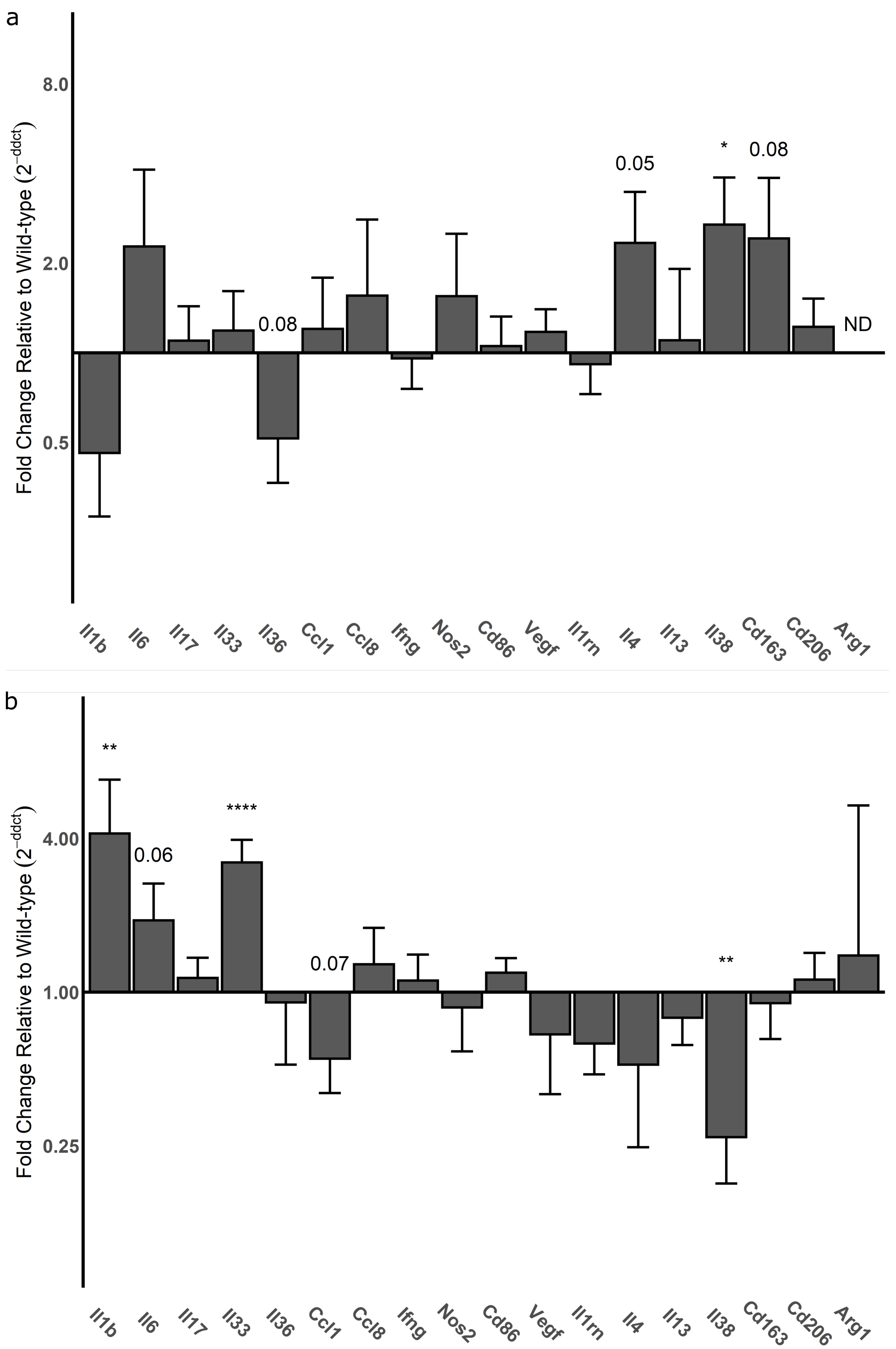


**Supplementary Figure 12: Screen of deficient versus wild-type gene expression in male mice.** qPCR results of pro-inflammatory versus pro-remodeling immune markers at day 3 **(a)** and day 11 **(b)** in male mice. Fold change and significance of deficient mouse sample gene expression was normalized relative to wild-type gene expression. Gene expression plotted as mean ± SEM. (ND = not detected, value displayed for trend $p \leq 0.1$, *$p < 0.05$, **$p < 0.01$, and ****$p < 0.0001$). n = 11-16 per group.

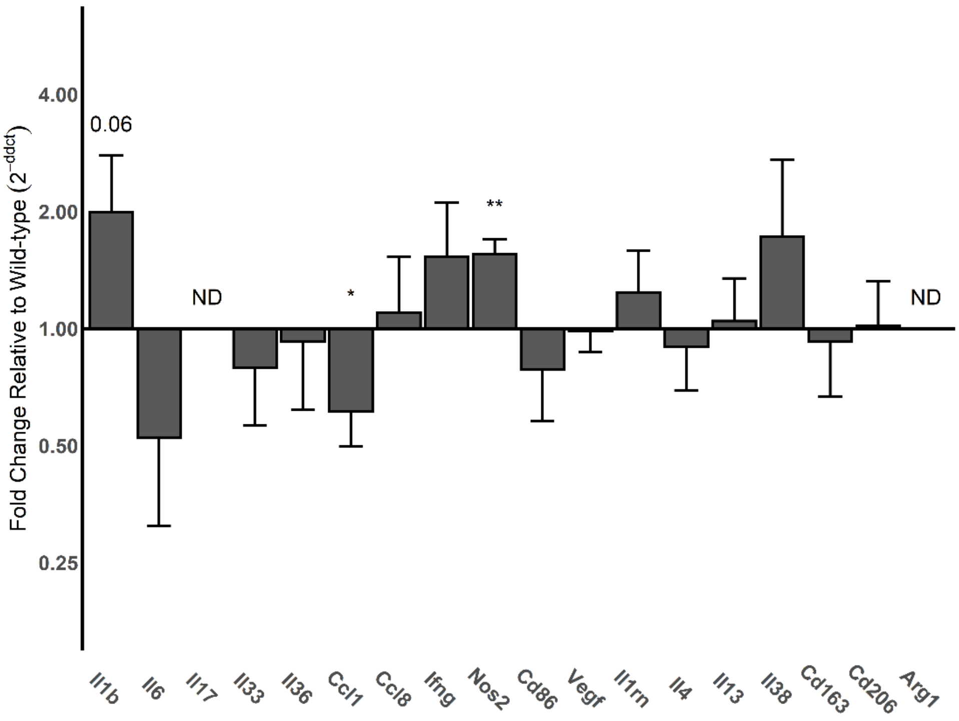


**Supplementary Figure 13: Screen of deficient versus wild-type gene expression in female mice.** qPCR results of pro-inflammatory versus pro-remodeling immune markers at day 3 in female mice. Fold change and significance of deficient mouse sample gene expression was normalized relative to wild-type gene expression. Gene expression plotted as mean ± SEM. (ND = not detected, value displayed for trend $p \leq 0.1$, *$p < 0.05$, **$p < 0.01$). n = 6-8 per group.

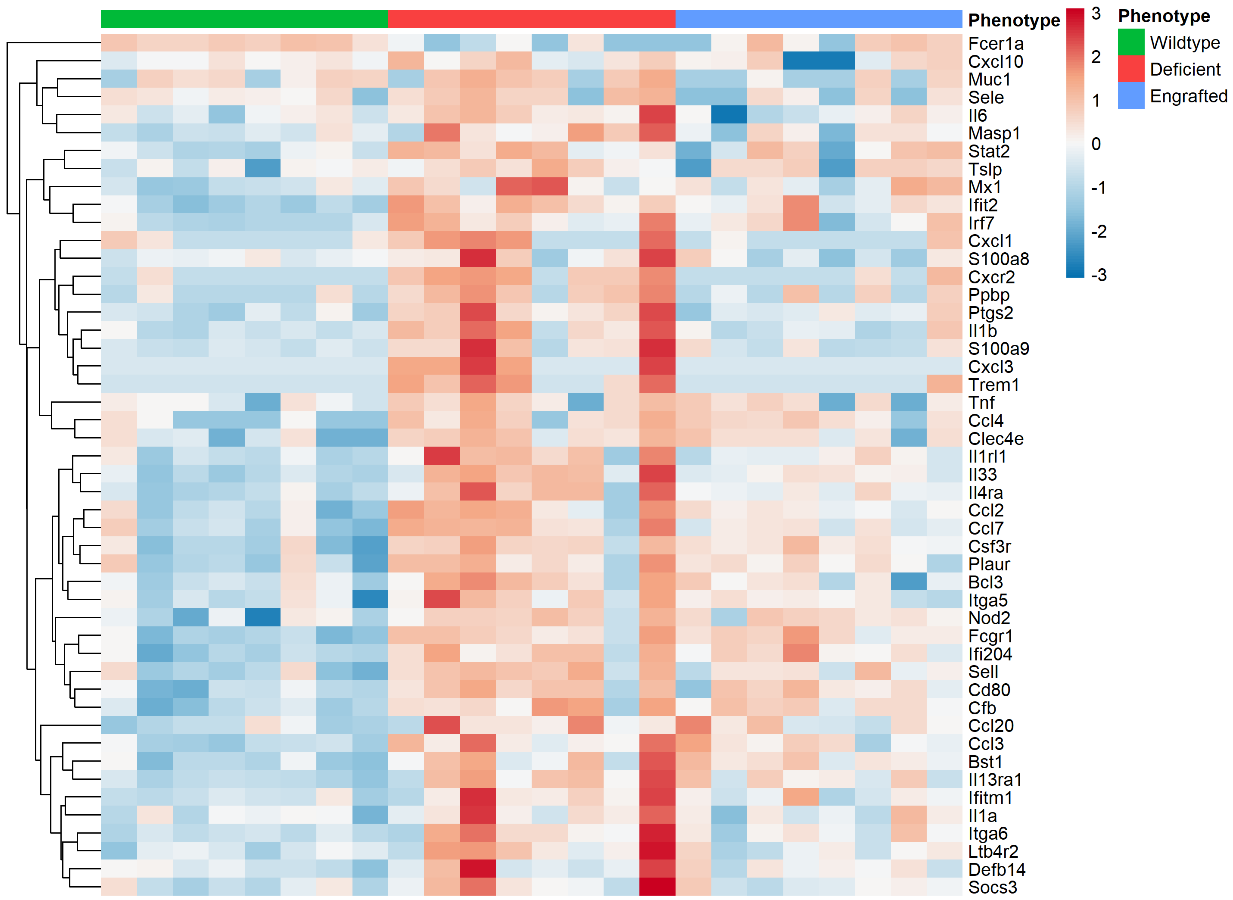


**Supplementary Figure 14: Heatmap of wild-type, deficient, engrafted gene expression**. Z-scores of differentially expressed genes based on q-value < 0.05 and log2 fold change ≥ 1. Nanostring nCounter gene expression data normalized based on the NanostringDiff package[2] method and displayed with pheatmap package[3] in R. Sample ID of wild-type (top green bar), deficient (top red bar), and engrafted mouse sample (top blue bar) for male mice at day 11 post-injection. Upregulated expression in red and downregulated in blue. n = 8 per group.

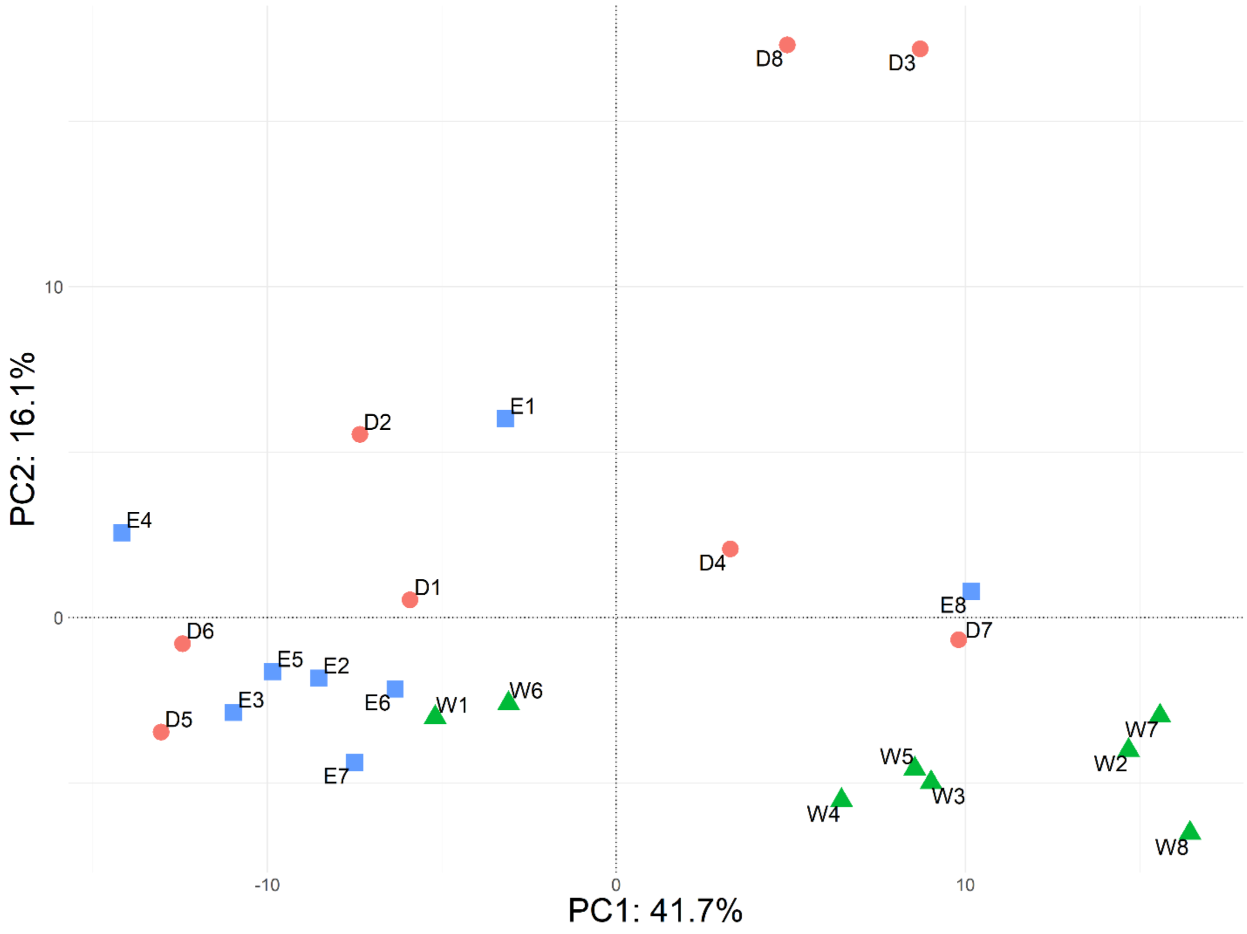


**Supplementary Figure 15: Principal component analysis plot of Nanostring nCounter data.** PC1 (41.7% of variance) versus PC2 (16.1% of variance) of the top 50% of assessed genes sorted by variance of their normalized gene expression based on the NanostringDiff package method. Points indicate wild-type, W1-W8 (green triangles), deficient, D1-D8 (red circles), and engrafted, E1-E8 (blue squares) samples. n = 8 per group.

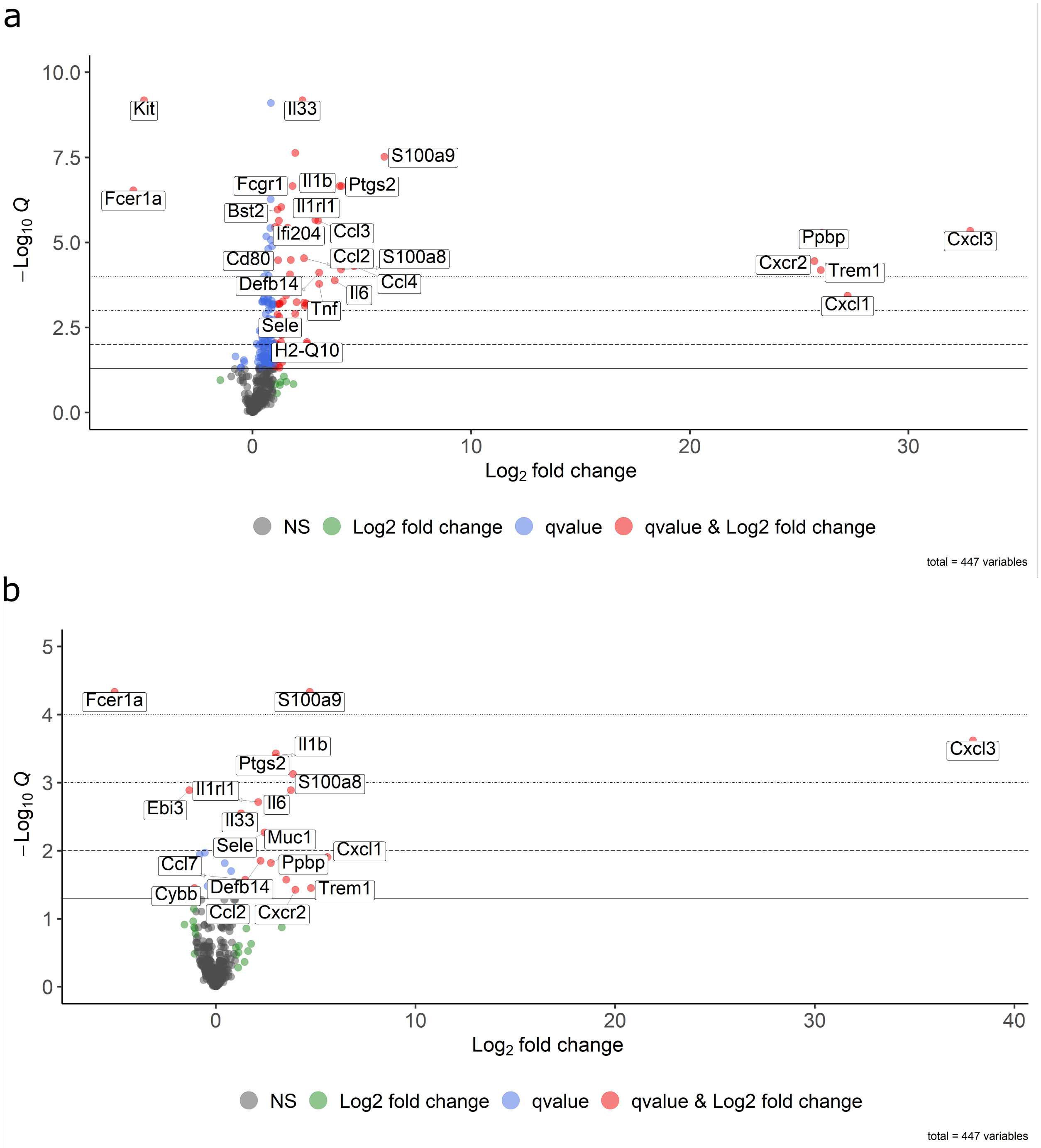


**Supplementary Figure 16: Volcano plots of pairwise comparisons for deficient response.** Top 25 ranked differentially expressed genes between deficient versus **(a)** wild-type and **(b)** engrafted sample groups with dots indicating false discovery rate less than 0.05 (blue, red), absolute fold change greater than 2 (green, red) or non-significant (black). Dashed lines indicating different significance thresholds. Results from n = 8 per group. (*$q$ < 0.05, **$q$ < 0.01, ***$q$ < 0.001 and ****$q$ < 0.0001).

**Table S1: Pro- and Anti-Inflammatory, and Pro-remodeling Immune Primers**

| Gene | | Forward Primer | | | Reverse Primer | |
|---|---|---|---|---|---|---|
| Arg1 | 5' | GAACACGGCAGTGGCTTTAAC | 3' | 5' | TGCTTAGTTCTGTCTGCTTTGC | 3' |
| Ccl1 | 5' | AGAAAGAGCTTCCCCTGAAGTTT | 3' | 5' | TGAGGCGCAGCTTTCTCTACC | 3' |
| Ccl8 | 5' | CCCCTTCGGGTGCTGAAAAG | 3' | 5' | TCACTGACCCACTTCTGTGTG | 3' |
| Cd163 | 5' | GAGACACACGGAGCCATCAA | 3' | 5' | CGTTAGTGACAGCAGAGGCA | 3' |
| Cd206 | 5' | GTGGACGCTCTAAGTGCCAT | 3' | 5' | GAATCTGACACCCAGCGGAA | 3' |
| Cd86 | 5' | GACTTGAACAACCAGACTCCTG | 3' | 5' | ATCAGCAAGACTGTCACAAAGA | 3' |
| Gapdh* | 5’ | CATCAAGAAGGTGGTGAAGC | 3’ | 5’ | GTTGTCATACCAGGAAATGAGC | 3’ |
| Ifng | 5’ | CGGCTGACTGAACTCAGATTG | 3’ | 5’ | CTGCAGCTCTGAATGTTTCTTAT | 3’ |
| Il13 | 5’ | GAGCAACATCACACAAGACCAGA | 3’ | 5’ | GGCCAGGTCCACACTCCATA | 3’ |
| Il17 | 5’ | AAGCTGGACCACCACATGAA | 3’ | 5’ | CCCTGAAAGTGAAGGGGCAG | 3’ |
| Il1b | 5’ | TGCCACCTTTTGACAGTGATG | 3’ | 5’ | TGATGTGCTGCTGCGAGATT | 3’ |
| Il1rn | 5’ | TCGGAGTACCTGTCATGCAAA | 3’ | 5’ | GCTTGCATCTTGCAGGGTCT | 3’ |
| Il33 | 5’ | TCCAACTCCAAGATTTCCCCG | 3’ | 5’ | CAGTGCAGTAGACATGGCAGAA | 3’ |
| Il36 | 5’ | AGGGCAAACCAACTTTGCAG | 3’ | 5’ | GAAGTGGAGCCCTCTATGCC | 3’ |
| Il38 | 5’ | TGCAGGAATGTGCTCCCTTC | 3’ | 5’ | GGTCTAGGCCTCGGTTAGGA | 3’ |
| Il4 | 5’ | GAGACTCTTTCGGGCTTTTCG | 3’ | 5’ | CAGTGATGTGGACTTGGACTC | 3’ |
| Il6 | 5’ | TCCAGTTGCCTTCTTGGGAC | 3’ | 5’ | AGTCTCCTCTCCGGACTTGT | 3’ |
| Nos2 | 5’ | CAGCTGGGCTGTACAAACCTT | 3’ | 5’ | CATTGGAAGTGAAGCGTTTCG | 3’ |
| Vegf | 5’ | GCACATAGAGAGAATGAGCTTCC | 3’ | 5’ | CTCCGCTCTGAACAAGGCT | 3’ |

*Housekeeping Gene

**Table S2: Estrogen Receptor Primers**

| Gene | | Forward Primer | | | Reverse Primer | |
|---|---|---|---|---|---|---|
| Esr1 | 5' | ACGCTCTGCCTTGATCACAC | 3' | 5' | CGAGTTACAGACTGGCTCCC | 3' |
| Esr2 | 5' | GGTCCTGTGAAGGATGTAAGGC | 3' | 5' | TAACACTTGCGAAGTCGGCAGG | 3' |
| Gapdh* | 5' | CCCCTTCGGGTGCTGAAAAG | 3' | 5' | TCACTGACCCACTTCTGTGTG | 3' |

*Housekeeping Gene
